# Stepwise Stiffening/Softening of and Cell Recovery from Reversibly Formulated Hydrogel Double Networks

**DOI:** 10.1101/2024.04.04.588191

**Authors:** Irina Kopyeva, Ethan C. Goldner, Jack W. Hoye, Shiyu Yang, Mary C. Regier, Kaitlyn R. Vera, Ross C. Bretherton, Cole A. DeForest

## Abstract

Biomechanical contributions of the ECM underpin cell growth and proliferation, differentiation, signal transduction, and other fate decisions. As such, biomaterials whose mechanics can be spatiotemporally altered – particularly in a reversible manner – are extremely valuable for studying these mechanobiological phenomena. Herein, we introduce a poly(ethylene glycol) (PEG)-based hydrogel model consisting of two interpenetrating step-growth networks that are independently formed via largely orthogonal bioorthogonal chemistries and sequentially degraded with distinct bacterial transpeptidases, affording reversibly tunable stiffness ranges that span healthy and diseased soft tissues (e.g., 500 Pa – 6 kPa) alongside terminal cell recovery for pooled and/or single-cell analysis in a near “biologically invisible” manner. Spatiotemporal control of gelation within the primary supporting network was achieved via mask-based and two-photon lithography; these stiffened patterned regions could be subsequently returned to the original soft state following sortase-based secondary network degradation. Using this approach, we investigated the effects of 4D-triggered network mechanical changes on human mesenchymal stem cell (hMSC) morphology and Hippo signaling, as well as Caco-2 colorectal cancer cell mechanomemory at the global transcriptome level via RNAseq. We expect this platform to be of broad utility for studying and directing mechanobiological phenomena, patterned cell fate, as well as disease resolution in softer matrices.

**TOC Description:** Biomaterials that can dynamically change stiffnesses are essential in further understanding the role of extracellular matrix mechanics. Using independently formulated and subsequently degradable interpenetrating hydrogel networks, we reversibly and spatiotemporally trigger stiffening/softening of cell-laden matrices. Terminal cell recovery for pooled and/or single-cell analysis is permitted in a near “biologically invisible” manner.

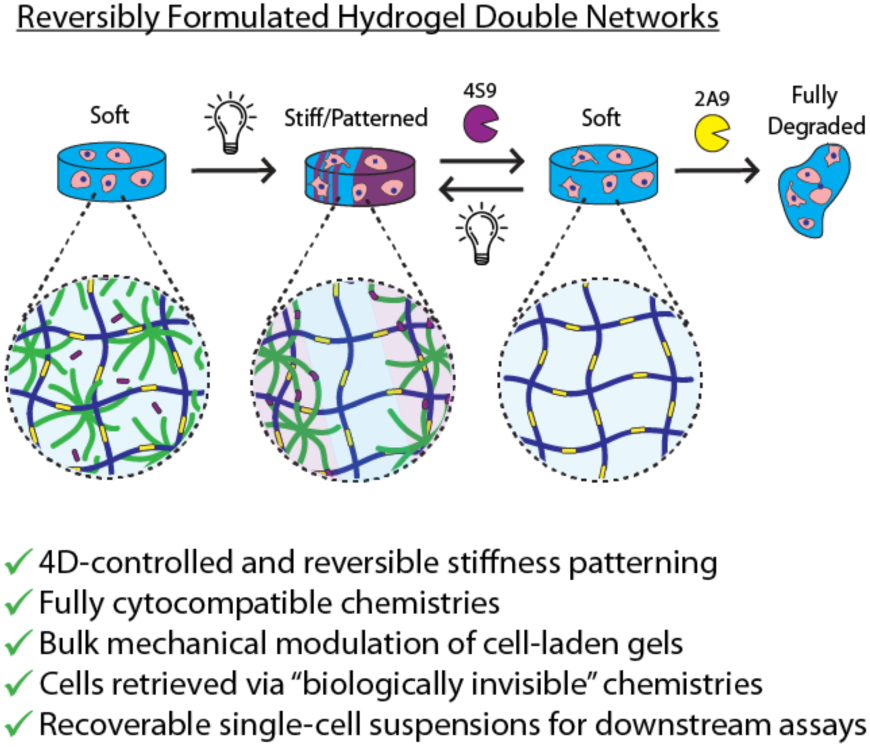

## Introduction

The interactions between cells and their environment are complex and dynamic. During development and disease, the extracellular matrix (ECM) undergoes major remodeling, resulting in changes in biochemical composition, biomechanical properties, and topography.^[1–3]^ ECM biomechanics are known to play a crucial role in cell growth, differentiation, proliferation, and signal transduction.^[4]^ The elastic modulus (E), or the stiffness, varies greatly among different tissues, ranging from 17 Pa (fat) to 20 GPa (cortical bone), supporting their diverging form and function.^[5]^ Cells translate mechanical stimuli in numerous ways; among the most studied mechanosensory complexes are focal adhesions, which initiate a complex signaling cascade when subjected to external force.^[4]^ Downstream, transcription factors such as Yes-associated protein (YAP) transmit cytoskeletal tension to the nucleus, impacting gene expression.^[6–8]^ Numerous diseases can be viewed in terms of mechanical dysregulation of tissue—two common examples being fibrosis and cancer. In fibrosis, fibroblasts respond to increased ECM stiffness and activate, secreting excess ECM proteins, which in turn further stiffens the matrix and perpetuates a self-amplifying feedback loop.^[9]^ In breast cancer and other solid tumors, lysyl-oxidase-mediated stiffening (from 150Pa to upwards of 6kPa) of the matrix drives malignancy.^[10,11]^

Synthetic hydrogels with viscoelastic and tunable properties akin to tissue offer an attractive platform to study mechanotransduction phenomena. Specifically, materials that can change their properties *in situ* afford researchers the ability to simulate the dynamic nature of the ECM.^[12]^ Stiffening hydrogels have allowed for the study of the initiation of fibrosis and mesenchymal and muscle stem cell differentiation.^[13–15]^ Yet, these systems cannot answer questions such as how cells respond to softening events, such as during cervical remodeling during pregnancy^[16]^ or whether pathologies, such as cancer progression can be reversed by modulating ECM mechanics.^[17,18]^ While photodegradable hydrogels have been developed for triggered material softening,^[19–23]^ these systems exhibit high light absorptivity/attenuation that confines softening to the surface; dynamic softening studies are thus limited to 2D-seeded cells. Other systems based on stimuli-responsive protein crosslinkers have achieved modulation of bulk mechanics, but do so with a limited dynamic range^[24]^ or have relied on chemistries that were not shown to be compatible with 3D cell encapsulation protocols.^[25,26]^ To improve upon these models, we aimed to design a dynamic hydrogel culture system whose mechanics could be reversibly patterned across a wide range of bulk stiffnesses, and whose encapsulated cells could be subsequently recovered in a “biologically invisible” manner for downstream pooled and/or single-cell-based analysis.

To address our design criteria, we turned to enzymatic degradation and identified the *S. aureus* transpeptidase, Sortase A (SrtA), as a bioorthogonal tool. Wild-type SrtA binds to the sorting motif “LPXTG” (where X is any amino acid) and cleaves between the Thr-Gly peptide bond. The Thr reacts with a Gly-Gly-Gly motif to form an LPXTGGG product, simultaneously displacing any sequence C-terminal to the sorting motif’s Thr. This sorting motif is extremely uncommon in the mammalian proteome, rendering this reaction largely bioorthogonal.^[27]^ SrtA has been utilized to rapidly release cells from hydrogels for downstream processing, as well as for modulating hydrogel mechanics, with no deleterious effects on survival and signaling.^[27–29]^ Wild-type SrtA has been evolved to yield two orthogonal sortases—eSrtA(2A9) and eSrtA(4S9) (respectively denoted 2A9 and 4S9) —that recognize with high specificity the sequences LAETG and LPESG, respectively.^[30]^ Previously, we have demonstrated that these peptide sequences can be included within single-network hydrogel crosslinkers to selectively release cells from materials in a “biologically invisible” manner, with minimal perturbation of the transcriptome of sensitive primary cell types; this method allowed for the construction of complex, multimaterial, 3D cell culture models using open-microfluidic patterning.^[28]^ Building on these efforts, we envisioned creating hydrogel double networks (DNs) – in which each interpenetrating single network could be independently formed via largely orthogonal bioorthogonal chemistries and subsequently degraded using orthogonal sortases – to permit step-wise stiffening, softening, and cell recovery via full network degradation (**Figure 1a**).

**Figure 1.**
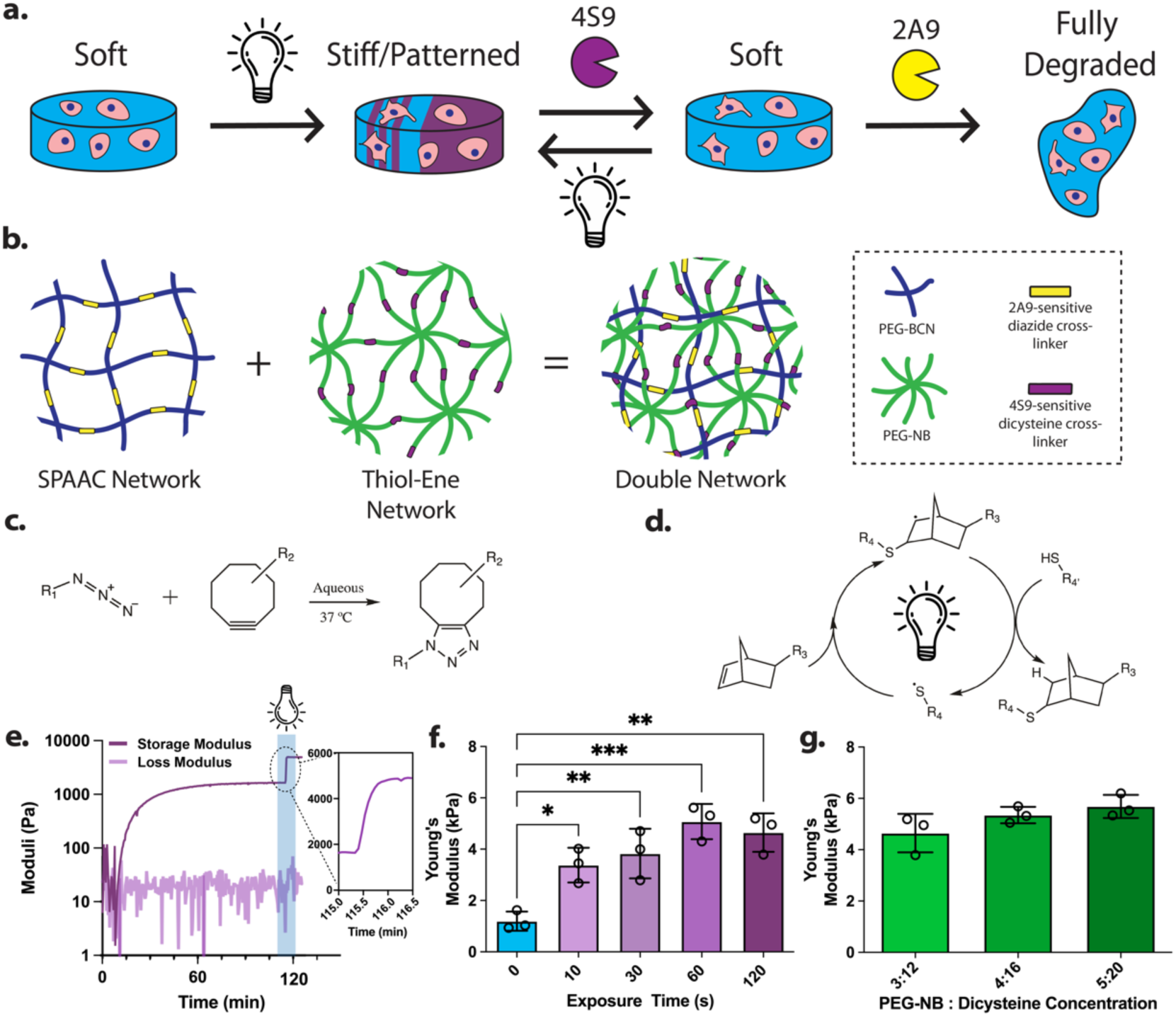
Double networks are tunably formed. **(a)** DNs are reversibly formulated with spatiotemporal control using a collection of orthogonal network formation and degradation chemistries. **(b-c)** DNs are composed of two distinct PEG-based networks formed via two popular bioorthogonal reactions: (b) strain-promoted azide-alkyne cycloaddition (SPAAC) and (c) the radical-mediated and light-driven thiol-ene reaction. **(d)** *In situ* photorheology demonstrates stepwise DN formation; SPAAC network formation proceeds spontaneously in the presence of thiol-ene network precursors, the latter of which is rapidly photopolymerized. Inset shows dramatic increase in storage modulus upon light exposure. **(e)** Young’s moduli as determined by AFM of swollen in PBS gels (initial concentrations 3 mM PEG-BCN: 6 mM diazide: 3 mM PEG-NB: 12 mM dicysteine: 1 mM LAP), post various light exposure times (0 – 2 min, 10 mW cm^-2^). SPAAC gels—1196 ± 370 Pa; 10 s—3381 ± 670Pa; 30 s—3829 ± 970 Pa; 1 min—5079 ± 690 Pa; 2 min—4648 ± 750Pa. One-Way ANOVA, Tukey’s post-hoc test. *p = 0.025, **p = 0.0078, ***p = 0. 0004. **(f)** Young’s moduli as determined by AFM of swollen gels, keeping SPAAC network constant, but varying molarity of thiol-ene network. 3mM PEG-NB—4648 ± 750 Pa; 4 mM PEG-NB—5352 ± 320 Pa; 5 mM PEG-NB—5687 ± 450 Pa.

Our DN unites two popular bioorthogonal reactions used for step-growth hydrogel biomaterial formation: a spontaneous strain-promoted azide-alkyne cycloaddition (SPAAC) between a terminal azide and a strained alkyne, and a thiol-ene photoreaction involving the radical-mediated addition of a thiol to a strained alkene (**Figures 1b,c**).^[31,32]^ Poly(ethylene glycol) (PEG) star polymers were end-functionalized with either bicyclononyne (BCN) or norbornene (NB) moieties, which respectively react with diazide- or dicysteine-containing peptides – each degradable with an orthogonal sortase and the same cell-secreted matrix metalloproteases (MMPs) – to form interpenetrating covalent networks. Notably, since SPAAC and thiol-ene reactions are kinetically orthogonal,^[33]^ DNs can be formed in a single reaction mixture via “self-sorting”.^[34]^ Further, the biochemical properties of each individual network can be distinctly tuned through pendant peptide incorporation (e.g., inclusion of an azide-modified RGDS peptide enables cell adhesivity to the SPAAC network, but not that formed via thiol-ene), a useful feature for maintaining constant material biofunctionalization throughout dynamic material stiffening/softening. We anticipate this platform will prove uniquely powerful towards studying mechanotransduction and memory in 4D.

## Materials and Methods

Complete experimental procedures are provided in the Supplementary Information, in particular for syntheses relating to macromer/peptide synthesis as well as sortase expression/purification that has been described previously.

### Materials

Materials, reagents, and cell culture consumables were purchased from Sigma-Aldrich (St. Louis, MO), ThermoFisher (Waltham, MA), and ChemImpex (Wood Dale, IL), unless otherwise noted. Fmoc-protected amino acids were purchased from ChemPep Inc. (Wellington, FL). Dicysteine peptide Ac-GCRDLPESGGPQGIWGQDRCG-NH_2_ (<u>4S9 degradable sequence</u>, <u>MMP-degradable sequence</u>) was purchased from Genscript (Piscataway, NJ), resuspended in 10% glacial acetic acid, and lyophilized to yield aliquots of the desired mass. Poly(ethylene glycol) tetra-bicyclononyne (PEG-BCN), 8-arm PEG-norbornene, and the diazide peptide crosslinker [N_3_-R<u>GPQGIWGQLAETG</u>GRK(N_3_)-NH_2_ (<u>2A9 degradable sequence</u>, <u>MMP-degradable sequence</u>)] were synthesized and processed as previously described (**Supplementary Methods S1-2**, **Supplementary Figure S1**).^[28,32,35,36]^ pET29b expression plasmids for 2A9 and 4S9 were a generous gift from Dr. David Liu at Harvard University (Addgene plasmids #75145 and #75146). Sortase enzymes were expressed and purified as previously described (**Supplementary Method S3**, **Supplementary Figure S2**).^[28]^ Caco-2 colorectal adenocarcinoma cells were a generous gift from Dr. William Grady at the Fred Hutch Cancer Center.

### Double Network Hydrogel Formation

DNs were formed in a stepwise manner. Unless otherwise specified, all monomers and gel precursors were combined at 3 mM PEG-BCN: 6 mM diazide: 3 mM PEG-NB: 12 mM dicysteine: 1 mM Lithium phenyl-2,4,6-trimethylbenzoylphosphinate (LAP; Allevi3D; Philadelphia, PA). Upon mixing, 10 µL droplets were pipetted between Rain-X®-coated glass slides spaced at 500 µm. SPAAC networks were allowed to form for 20 minutes at 37 °C, at which point they were exposed to collimated near-UV light (λ = 365 nm; 10 mW cm^-2^; OmniCure 1500) for 2 minutes (unless otherwise specified) to drive thiol-ene polymerization. For encapsulation, gel formulations included 1 mM N_3_-GRGDS-NH_2_ to promote cell adhesion.

### Photorheology

Gel formation kinetics and storage moduli (G’) were analyzed on a Physica MCR-301 rheometer (Anton Paar; Graz, AT) at 25 °C with 8 mm parallel plate geometry (0.5 mm gap, 1Hz, 1% strain). The SPAAC network was allowed to form for 1 hr, at which point 365 nm light at 10 mW cm^-2^ was irradiated from the bottom plate coupled to a fiber optic light guide from a multiwavelength LED light source (Mightex Systems; Toronto, ON). The frequency and strain were determined to fall within the viscoelastic range via frequency and amplitude sweeps.

### Atomic Force Microscopy Measurements

40 µL of hydrogel solution was pipetted onto a Rain-X®-coated glass slide with 500 µm-thick rubber spacers, on top of which were placed 18 mm-diameter thiolated glass coverslips. After gel formation, coverslips with attached gels were allowed to swell overnight in PBS. The following day, atomic force microscopy (AFM) measurements were performed on an Asylum Cypher S AFM (Oxford Instruments; Concord, MA) with a Bruker NP-O10 silver nitride cantilever (Bruker; Camarillo, CA) (k = 0.35 N/m; f = 65 kHz) functionalized with 50 µm-diameter soda lime glass spherical beads (Cospheric; Santa Barbara, CA).^[37]^ The sensitivity and the spring constant of the probe was calibrated before usage (k = 0.22 N/m) in PBS using a fused silica sample in a Peak Force QNM Sample kit (Bruker; Camarillo, CA). All measurements were taken in contact mode and approach and retraction speeds were 1.98 µm s^-1^ with a trigger point of 1V and a retraction distance of 1 µm. For each sample, 5 force maps were generated by indenting in 30 µm x 30 µm grids with 100 indentation points, collecting at least 300 fittable force curves. Three substrates were measured for each condition. To evaluate Young’s moduli, indentations were fitted to a Hertzian model using the Cypher 16.14.216 software. For softened hydrogels, due to the heterogenous surface heights, samples were indented in 5 discrete locations at least 10 times, giving at least 50 measurements for subsequent analysis per substrate. For patterned hydrogels, the same procedure was used as for softened hydrogels, except that 5 discrete locations were chosen in the light-exposed regions (“in” pattern) and the non-exposed region (“out” of pattern). Three samples were measured for each condition.

### Double Network Softening

DNs were softened by treating with 50 µM 4S9, 18 mM GGG, and 10 mM CaCl_2_ for 1 hour, and washed 3 x 10 mins afterwards to remove degraded thiol-ene network components. For cell culture applications, GGG was resuspended in full-serum media and the pH was adjusted to ∼7; additionally, all components were sterile filtered through 0.2 µm syringe filters. Post-softening, gels were washed 3 x 10 mins with full-serum media.

### Assaying Orthogonal Degradation of DNs

Hydrogels were made at the following concentrations: 3 mM PEG-BCN: 6 mM diazide: 3 mM PEG-NB: 12 mM dicysteine: 1 mM LAP. Each network was tagged with a different fluorophore: dicysteines resuspended at a 40 mM concentration were prereacted with 100 µM of AlexaFluor 488 maleimide (1:400 dye:peptide) (Click Chemistry Tools; Scottsdale, AZ) for 15 minutes, whereas PEG-BCN was preincubated for 15 minutes with 50 µM of AlexaFluor 568 azide (1:200 dye:PEG) (Click Chemistry Tools; Scottsdale, AZ). DN hydrogels were then formed with these fluorescently tagged components, and allowed to swell and wash away unreacted dye overnight. The following day, gels were treated with 50 µM of either 2A9 or 4S9, 18 mM GGG, and 10 mM CaCl_2_ (400 µL total volume). To quantify degradation extent, 2 µL of supernatant was taken from the well at each timepoint and diluted in 98 µL of PBS in a black 96-well plate. Fluorescent values were read on a plate reader (Molecular Devices; San Jose, CA) and normalized to the final release at 12-hour. To normalize the non-degrading network, the gels were treated with the other sortase variant for 1 hour at the end of the experiment to fully degrade the remaining network (e.g., a DN analyzed for 2A9 degradation was treated at the end with 4S9 to degrade the remaining single network).

### Double Network Patterning and Visualization

DNs were formed as described previously at 3 mM PEG-BCN: 6 mM diazide: 3 mM PEG-NB: 12 mM dicysteine: 1 mM LAP. 100 µM FAM- maleimide was additionally included to visualize the thiol-ene network. 10 µL gels were formed on Rain-X^®^-coated glass slides with 1 mm-thick rubber gaskets. Gels were incubated at 37 °C for 20 minutes to form initial SPAAC networks. After initial polymerization, one glass slide was removed and replaced with a photomask to reduce diffraction during patterning. Gels were then exposed to collimated UV light (λ = 365 nm; 10 mW cm^-2^; Omnicure 1500) through a chrome photomask (Photo Sciences; Valencia, CA) for 2 minutes, allowing for thiol-ene polymerization and formation of DN patterns. Following patterning, gels were soaked in 10 µM Cy5-N_3_ (Click Chemistry Tools; Scottsdale, AZ) for 2 hours at 37 °C to label the SPAAC network and then washed 3 x 1 hr in PBS. For cellular applications, the SPAAC network was not fluorescently labelled, and gel formulations included 1 mM N_3_-GRGDS-NH_2_ to promote cell adhesion. Imaging was performed using the Leica Stellaris confocal microscope.

### Two-photon Patterning and Visualization

DNs were formed as described previously (3 mM PEG-BCN: 6 mM diazide: 3 mM PEG-NB: 12 mM dicysteine: 1 mM LAP), but with 25 µM Rhodamine included as a photosensitizer. Gels were formed between Rain-X®-coated glass coverslips and slides with 1 mm-thick rubber gaskets that enclosed the gel in an air chamber. Gels were patterned using a Thorlabs Bergamo II 2-photon confocal microscope equipped with a Coherent Chameleon Discovery NX tunable femtosecond laser and an Olympus liquid-immersion objective (25X, numerical aperture = 0.95), controlled via ScanImage software (MBF Bioscience; Williston, VT).^[38]^ The imported image sequence (field of view = 450 µm × 450 µm × 313 µm) was patterned at 750 nm in the z-direction (resolution = 0.44 µm/px × 0.44 µm/px × 1 µm/px, pixel dwell time = 3.2 µs) (**Supplementary Figure S3**). Each image within the sequence was repeated 50 times at 270 mW. Following patterning, gels were soaked in 10 µM Cy5-N_3_ (Click Chemistry Tools; Scottsdale, AZ) for 2 hours at 37 °C to block any unreacted BCN groups, washed 3 x 1 hr in PBS, and then incubated with 50 µM AZDye 488-tetrazine (Click Chemistry Tools; Scottsdale, AZ) for 2 hours at 37 °C to visualize the thiol-ene network. Gels were washed overnight in PBS and then imaged on a Leica Stellaris confocal microscope.

### Cell Culture and Encapsulation

10T1/2 fibroblasts were cultured in Dulbecco’s Modified Eagle Medium (DMEM) supplemented with 10% fetal bovine serum (FBS) and 1X Penicillin-Streptomycin (PS) in a standard 37 °C, 5% CO_2_ cell culture incubator. Cells were passaged 1:10 upon reaching 80% confluency. Experiments were conducted with 10T1/2 cells below passage 15. Human mesenchymal stem cells (hMSC) were purchased from RoosterBio, Inc. (Frederick, MD) and grown in DMEM supplemented with 10% FBS and 1X PS, and passaged similarly to 10T1/2 cells; all experiments were conducted with cells below passage 5. Caco-2 cells were cultured in Eagle’s Minimum Essential Medium (EMEM) supplemented with 20% FBS and 1X PS, and passaged similarly to the other cells; all experiments were conducted with cells below passage 5.

### Live/Dead Imaging

10T1/2 fibroblasts were encapsulated in gels at a concentration of 2 x 10^6^ cells mL^-1^ and cultured for 7 days. Dynamic DNs were treated on day 3 with 4S9. Cell viability was assayed by live/dead staining with calcein AM and ethidium homodimer (EtHD). Hydrogels were incubated in live/dead staining solution (2 µM calcein and 4 µM EtHD in PBS) for 1 hour prior to confocal imaging. Live/dead count was quantified from 250 µm max intensity projections (MIP) using IMARIS.

### Actin Staining and Analysis

hMSCs were encapsulated in gels at a concentration of 2 x 10^6^ cells mL^-1^ and cultured for 7 days in either bulk or patterned gels. On day 7, gels were fixed by treatment with 4% paraformaldehyde for 1 hr at room temperature, washed 3 x 10 mins with tris-buffered saline (TBS), and permeabilized for 30 mins with 0.5% Triton X-100 in TBS. Subsequently, actin was labelled with 1:400 Phallodin-532, and nuclei — with 1:1000 Hoechst 33342 in TBS. Gels were rinsed in TBS and imaged on a Leica Stellaris confocal microscope. Cell area and eccentricity were analyzed from 100 µm MIPs with Cell Profiler 4.0.^[39]^

### Lentiviral Construct Assembly and Transduction

YAP/TAZ-TRE-mStrawberry reporter was a gift from Ravid Straussman (Addgene plasmid # 158682). From this plasmid, polymerase chain reaction (PCR) was used to amplify the backbone and the TEAD-binding sequence repeats, TRE/GTIIC. The resultant PCR product was reassembled into a tdTomato-containing vector via Gibson assembly (NEB) with a tdTomato insert (sequence ordered as a codon-optimized gBlock from IDT) and transformed into NEB Stable *E. Coli*.

Human embryonic kidney (HEK293T) cells were plated at ∼40% confluency and allowed to adhere to tissue culture plastic overnight. Fresh media was added, and then HEKs were transfected with envelope plasmid pMD2.G (Addgene #12259), packaging plasmids pMDLG/pRRE (Addgene #12251) and pRSV-REV (Addgene #12253), and the YAP/TAZ-TRE-tdTomato plasmid using Lipofectamine 2000 (Invitrogen). Cells were cultured for 2 days post-transfection, and virus-laden media was harvested. Viral media was filtered (0.45 μm) and tested using lentiviral titration card (ABM Biologics). Active lentivirus was concentrated by mixing viral media with 4X lentiviral concentration solution (40% w/v PEG-8000, 1.2 M NaCl), vigorously shaking for 60 seconds, and agitated overnight at 4 °C. The following day, flocculated lentiviral particles were pelleted at 1600×g for 60 minutes at 4 °C, and supernatant was aspirated. Pellet was resuspended at 10X relative to initial viral media volume in PBS.

hMSCs were plated at ∼40-60% confluence the day before the transduction and allowed to adhere overnight. The following day, their media was replenished and supplemented with 8 μg mL^-1^ polybrene and the prepared lentivirus. Transduction was allowed to occur overnight, then viral media was aspirated and replaced with regular culture media. After 2 days in recovery, hMSCs were exposed to 4 μg mL^-1^ of blasticidin and were selected for 7 days. Selection media was replaced every 3-4 days. After selection, hMSCs were expanded and used for encapsulation.

For live-cell imaging of TEAD-binding activity, transduced hMSCs were encapsulated at a concentration of 2 x 10^6^ cells mL^-1^ and cultured for 7 days. Throughout the experiment, cells were live imaged on days 1, 3, 5, and 7 with 50 µm thick z=stacks in 3 discrete location per gel for quantification of tdTomato intensity with CellProfiler 4.0. Dynamic gels were treated with 4S9 immediately post imaging on their respective days (e.g., dynamic day 3 gels were first imaged on day 3, then softened). Following imaging on day 7, gels were fixed and labelled with phalloidin- 532 and Hoechst 33342.

### Bulk RNA Sequencing

Caco-2 cells were encapsulated in gels and cultured for 7 days, with a subset of DNs being softened on days 3 and 5. To induce full gel dissolution at the end of the tissue culture period, all gels were treated with 50 µM 4S9, 50 µM 2A9, 36 mM GGG, and 1 mM CaCl_2_ for 30 minutes. Cells were pelleted, washed 1X in ice-cold PBS, and lysed in TRIZol reagent. RNA was isolated from lysed samples using the Direct-Zol RNA microprep kit (Zymo Research; Irvine, CA), snap frozen in liquid nitrogen, and stored at -80 °C until further use. Prior to sequencing, RNA integrity (RIN) was evaluated using the 4200 TapeStation System (Agilent; Santa Clara, CA); all RIN values were greater than 8. mRNA was converted into dual-indexed cDNA libraries using the Illumina Stranded mRNA Prep and Ligation kit (Illumina; San Diego, CA) and sequenced on the Illumina NextSeq 2000 Platform (30 million clean paired-end reads per sample). Data analysis was conducted on the Partek Flow Software (Partek; St. Louis, MO). Reads were aligned using the STAR package,^[40]^ annotated to the human genome (Ensembl Transcripts release 109), and evaluated for differential expression via DESeq2.^[41]^ A false discovery rate (FDR) of 0.05 was used as a cut-off for significant differences in expression. Gene set enrichment analysis was performed using the Molecular Signatures Database (MSigDB).^[42]^

### Statistical Analysis

Unless otherwise noted, data and statistical analysis were conducted on GraphPad Prism 7.0.

## Results

### Double Network mechanical properties are tunable

We began by establishing that our baseline DN could be formed in a one-pot, stepwise manner via photorheology. We allowed the primary SPAAC network to first form over the course of an hour and then illuminated the single-network hydrogel with near-UV light (λ = 365 nm, 10 mW cm^-2^, 2 minutes) to form the interpenetrating secondary thiol-ene network. The storage modulus of the SPAAC network plateaued to 2241 ± 670 Pa at ∼1 hour, and then sharply increased to 4848 ± 890 Pa upon photopolymerization of the thiol-ene secondary network (**Figure 1d**). Next, having established that our two-step method would yield the expected step up in shear storage modulus, we explored varying parameters to create DN materials of various initial stiffnesses. Increasing the exposure time (0 – 2 min) led to significant increases in the Young’s moduli of swollen gels, as measured by atomic force microscopy (AFM) (**Figure 1e**); however, we observed a plateau after 1 minute of exposure time, indicating that the thiol-ene polymerization completed in that time. Similarly, increasing the final concentration of the thiol-ene network in the gel formulation led to increases in swollen Young’s moduli with the maximal 5 mM PEG-NB: 20 mM dicysteine condition achieving a modulus of 5687 ± 260 Pa (**Figure 1f**); however, these differences were not statistically significant. Additionally, we tested Young’s moduli of gel concentrations used in subsequent cell encapsulation experiments with and without N_3_-RGDS-NH_2_ to facilitate adhesion to the matrix,^[28]^ and noted a small and expected, but nonsignificant, decrease in Young’s modulus when RGDS was included (**Supplementary Figure S4**).

### Sortase treatments proceed orthogonally and yield a step down in Young’s modulus

Our next goal was to demonstrate that our proposed method would allow for bulk hydrogel softening, achieved through conversion of our DN gels to a single network through sortase-mediated degradation. While either network can be crosslinked with either eSrtA-sensitive crosslink, for this initial study, we chose to include the 2A9-sensitive sequence “LAETG” on the diazide crosslinkers in the SPAAC network, and the 4S9-sensitive sequence “LPESG” on the dicysteine crosslinkers in the thiol-ene network (**Figures 2a-b**). Each peptide crosslinker also included the MMP-sensitive sequence “GPQGIWGQ” to allow for cell-mediated remodeling of the hydrogel matrix.^[36,43]^

**Figure 2.**
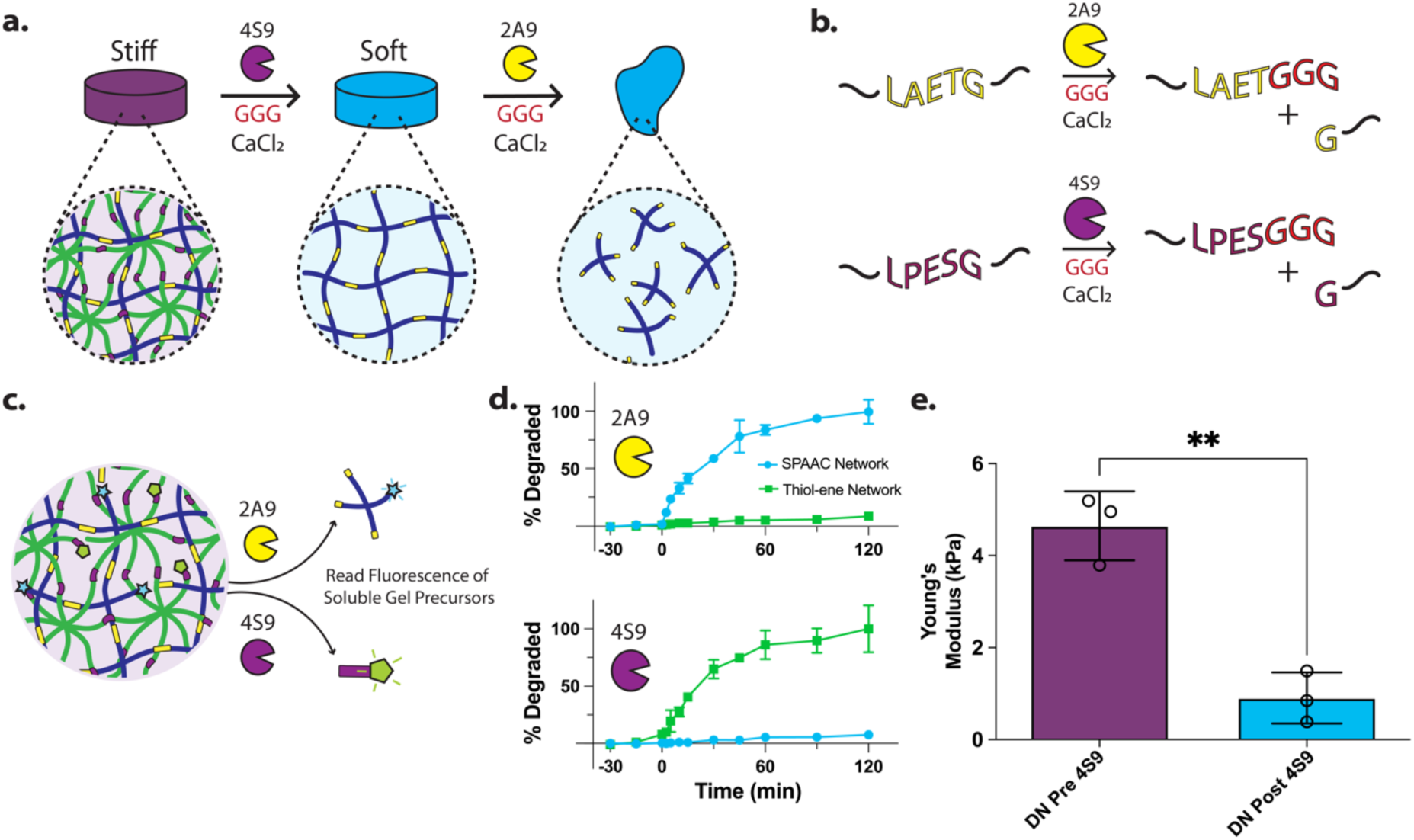
Double networks can be independently degraded in a stepwise manner. **(a)** Double networks are first treated with 4S9 to remove the thiol-ene network, and then fully degraded by treatment with 2A9 to yield fully soluble macromolecular building blocks. **(b)** Peptide recognition sequences for 2A9 and 4S9 included in hydrogel crosslinkers and degradation reaction post sortase treatment. **(c)** Schematic depicting individual labeling of each network with distinct fluorophores, and the monomeric component released upon each sortase treatment, tracked by increases in supernatant fluorescence. **(d)** Fluorophore release studies. At time = 0 min, 18 mM GGG, the respective sortase, and 1 mM CaCl_2_ were added to the solution the DN hydrogels were in. Hydrogel degradation was tracked by monitoring supernatant fluorescence, with values normalized to those obtained from 100% degraded gels 12 hours post reaction. **(e)** AFM measurements of DN gels pre- and post-4S9 treatment. DN pre 4S9 treatment: 4648 ± 750 Pa; DN post 4S9 treatment: 908 ± 550 Pa. Unpaired t-test, **p = 0.0024.

While orthogonal sortase variants have been successfully deployed for degrading complex layered hydrogels,^[28]^ this method has yet to be applied to selectively degrading interpenetrating networks, although other nonbioorthogonal enzymes (e.g. alginate lyase, hyaluronidase) have previously been employed;^[6,44]^ in fact, to our knowledge, there is not a prior report of using any stimuli to convert a synthetic polymer-based DN to a single network system. To monitor degradation via changes in supernatant fluorescence, we created DNs whereby each network was sparsely tagged with a different fluorophore (**Figure 2c**), which upon cleavage and release, diffuse into the supernatant. Whereas others have previously pre-incubated gels with sortase to allow for maximal diffusion and fast degradation kinetics,^[27,28]^ given our ultimate need to conduct such degradation in the presence of cells, we chose to add both sortase and GGG simultaneously. Upon the addition of 50 μM sortase solution, 18 mM GGG, and 10 mM CaCl_2_ at t = 0, we observed rapid degradation. Treatment with 2A9 led to degradation of the SPAAC network, with very minimal perturbation of the thiol-ene network, as expected, and conversely, treatment with 4S9 led to degradation of the thiol-ene network, with minimal impact on SPAAC network integrity (**Figure 2d**). Our results coincided with our previous findings, where we demonstrated that 2A9 and 4S9 can degrade multimaterials in a fully orthogonal manner.^[28]^ Intriguingly, degradation times for a 4-arm SPAAC network and an 8-arm thiol-ene network were almost identical: both reactions neared completion in 60 minutes. To maximize the mechanical difference between stiff and soft regimes, we chose to first treat our DNs with 4S9 to degrade the 8-arm thiol-ene network (**Figure 2a**). AFM measurements pre- and post-4S9 treatment similarly demonstrated a significant drop in Young’s modulus, with the softened modulus (878 ± 350 Pa) not statistically different from a single network SPAAC gel (1196 ± 210 Pa) (**Figure 2e**). The initial DN moduli (4.6 – 5.7 kPa) presented match that of softer tissues, including that of diseased kidney glomeruli,^[45]^ metastatic and primary colon cancer,^[17,46]^ pancreatic ductal adenocarcinoma,^[47]^ as well as other healthy examples.^[48]^ Softened moduli, similarly, match that of kidney, colon, and pancreatic healthy tissue.^[48]^

### Hydrogel formulations and 4S9 treatment are cytocompatible

After having established the ability to modulate bulk mechanics, we next assessed the suitability of this platform and method for 3D cell culture. 10T1/2 murine fibroblasts were encapsulated in SPAAC, thiol-ene, or DN gels and cultured for 7 days. A subset of the DNs were treated with 4S9 on day 3 of culture (“dynamic” condition) to soften the gel substrate and then cultured until the terminal timepoint of day 7. All gel conditions displayed a statistically indistinguishable and high viability (>91%) (**Figures 3a-b**), indicating that all gelation and softening conditions were mild and cytocompatible, in accordance with previous reports.^[27–29,49,50]^

**Figure 3.**
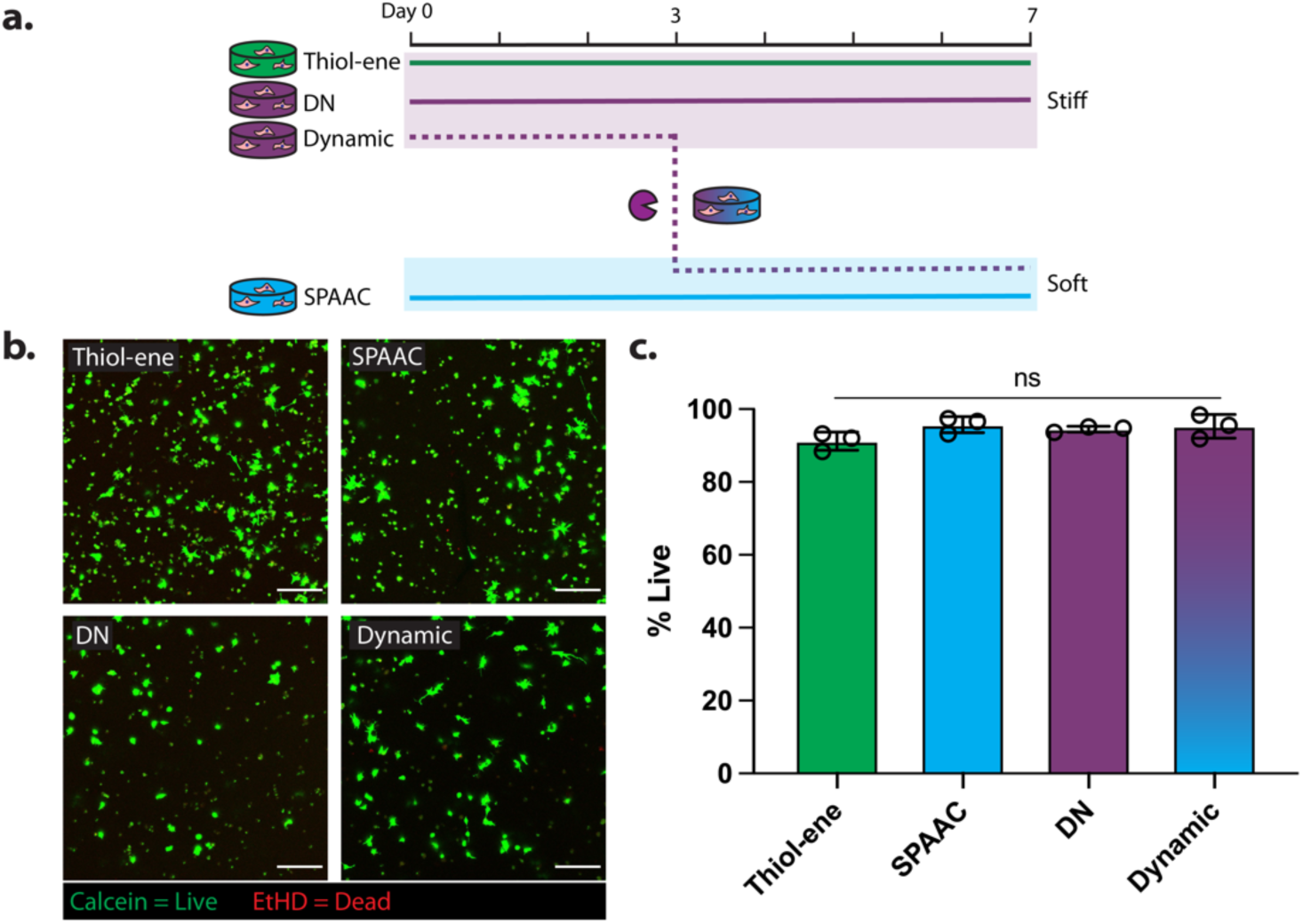
DNs can be formed and dynamically softened in a cytocompatible manner. **(a)** Experimental set-up for viability measurements. Static controls of thiol-ene, DN, and SPAAC gels were compared against dynamic DN gels treated with 4S9 on day 3 of culture. **(b)** Maximum Image Projection (MIP) of representative images (z = 250 µm). Live/Dead staining of encapsulated 10T1/2 fibroblasts shows excellent cytocompatibility of all possible network types on day 7 of culture. Scale bar = 100 µm. **(c)** Quantification of viability.

### DN formation enables spatiotemporal control over mechanical properties

Sortase-mediated reversible modulation of mechanics has been reported previously;^[29]^ however, prior reports have not demonstrated spatial control over mechanical properties, nor complete cell recovery following 3D culture. We leveraged the DN design to spatially control thiol-ene polymerization via mask-based lithography, yielding highly defined mechanical patterns that could then be subsequently removed using 4S9 (**Figure 4a-b**). To measure Young’s moduli prior to and post-4S9 treatment, gels were covered with a rough photomask hiding half of the gel and exposed to light; AFM indentation measurements were then taken on both halves of the gel. Subsequently, the gels were then treated with 4S9 and again, Young’s moduli measurements were taken on both sides. Prior to 4S9 treatment, the exposed regions exhibited Young’s moduli of 3258 ± 510 Pa, whereas the nonexposed regions were 1061 ± 320 Pa, and post treatment the previously patterned regions dropped to 838 ± 54 Pa, while the nonexposed regions remained at 1048 ± 490 Pa, which were not statistically different between each other (**Figure 4c**). While our chosen gelation chemistries permit one-pot formation of a DN, an alternative strategy is to diffuse in photocrosslinkable monomers into a pre-formed hydrogel prior to photostiffening.^[51]^ With this in mind, we sought to reversibly cycle between a single network hydrogel and a spatially patterned DN system. 24 hours post SPAAC gel formation, we incubated the gels in thiol-ene components for 4 hours at 37 °C, and then selectively exposed the gel to photomasked light to yield well-defined patterned thiol-ene polymerization (**Figure 4d**). The next day, the patterns were removed via 4S9 treatment, converting the DN back to a single network with no thiol-ene network remaining. We then repeated the patterning process, converting our single network SPAAC gel back into a DN system via thiol-ene polymerization, which proved successful. This process is likely repeatable many more times over, though extended diffusion times place practical limits on network cyclability.

**Figure 4.**
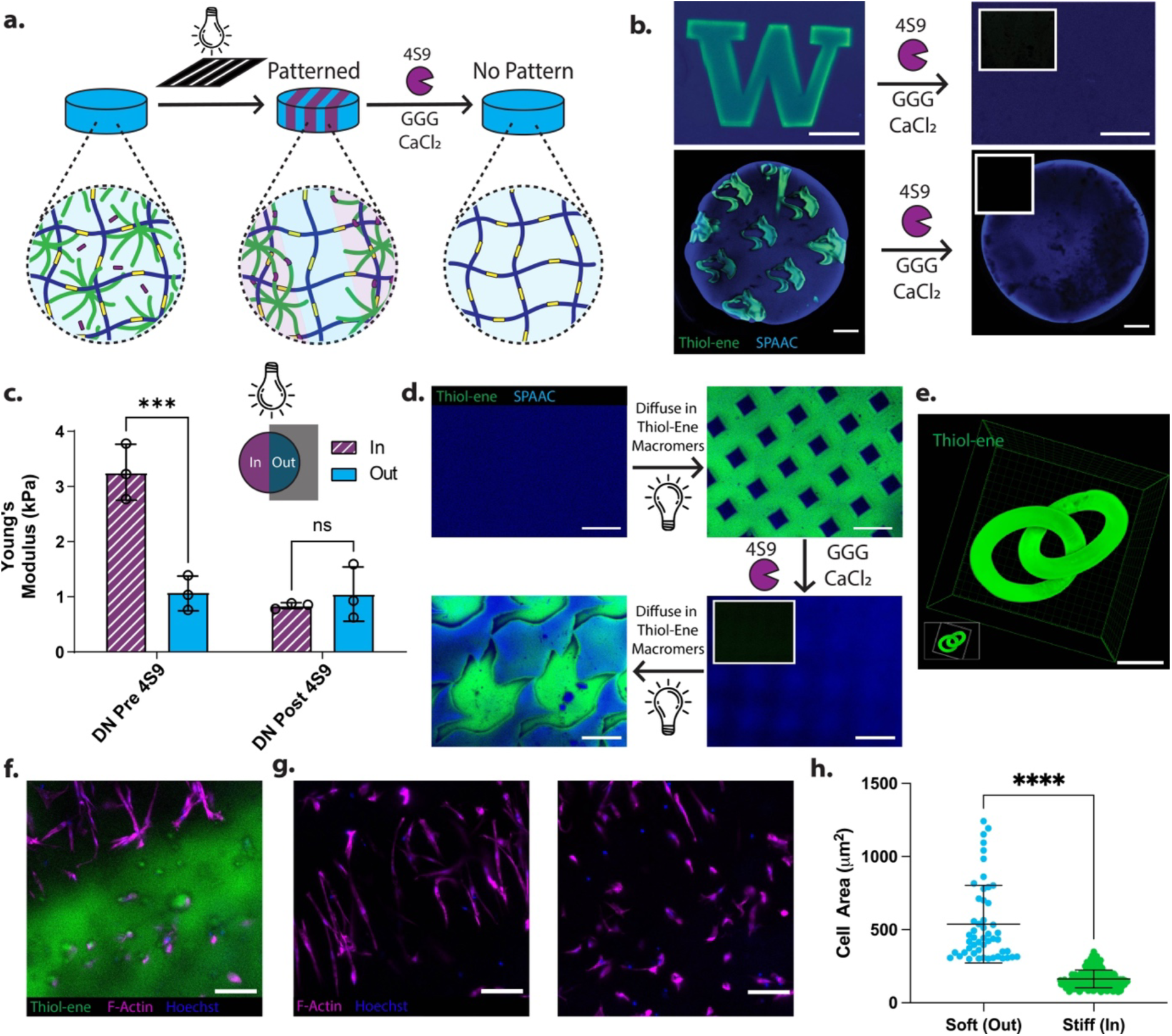
Double networks can be reversibly and spatiotemporally patterned to drive changes in encapsulated cell morphology. **(a)** Schematic depicting stepwise patterning and pattern removal. Soluble monomeric precursors can be mixed together in a one-pot mixture. SPAAC stepwise network formation occurs spontaneously, while thiol-ene polymerization can be spatially controlled photolithographically. Subsequently, thiol-ene patterns are removed with sortase 4S9 treatment. **(b)** Stiff patterns in a bulk hydrogel are enabled by localized thiol-ene polymerization and can be removed by 4S9 treatment. Insets depict no fluorescence is visible in the FAM channel (thiol-ene network) post enzymatic treatment. Top scale bar = 200 µm, bottom scale bar = 1 mm. **(c)** AFM measurements of half-patterned gels. “In” denotes a stiff region exposed to light, whereas “out” denotes the covered, non-exposed region. Two-Way ANOVA, ***p = 0.0002. **(d)** DN design allows for reversible patterning of mechanics. Thiol-ene gel components can be diffused into single network at later time points for mechanical patterning and can be reversibly removed and reinstated by rounds of 4S9 degradation and photopolymerization. Scale bar = 250 µm. **(e)** Intricate DN formations can be patterned using multiphoton laser-scanning lithography. Scale bar = 100 µm. **(f)** hMSCs encapsulated in stiffness-patterned hydrogels. Image shows the interface of stiff and soft regions. Scale bar = 100 µm. **(g)** hMSCs in soft (left) vs stiff (right) regions of patterned hydrogel. **(h)** Quantification of cell area in soft and stiff regions. Unpaired t-test, ****p < 0.0001.

Previous reports indicated LAP to be sensitive to two-photon-based activation;^[52–54]^ thus, we hypothesized that our DN system could enable the creation of complex 3D thiol-ene structures supported by the surrounding SPAAC matrix. Since LAP is only weakly two-photon active, we utilized rhodamine as a photosensitizer in conjunction with the photoinitiator to photopolymerize the second network in a set of intertwining rings via laser-scanning multiphoton lithography (**Figure 4e, Supplementary Figure S3**). We note that such complexity in patterned stiffness is currently not readily obtained via other additive methods, including conventional digital light projection, extrusion-based, and volumetric 3D printing.

Next, we asked whether these differences in patterned gel stiffness could elicit a change in cellular morphology. Human mesenchymal stem cells (hMSCs) were encapsulated in DNs featuring 200 µm-wide line patterns and cultured for 7 days, at which point they were fixed and stained with phalloidin to visualize cytoskeletal area (**Figures 4f-g**). We observed that cells encapsulated in the softer SPAAC regions (“out” of pattern) exhibited a significantly larger surface area compared to those in the stiffer DN regions (“in” pattern) (**Figures 4g, h**). We did note a surprising alignment of cells perpendicular to the pattern. We attribute this to differential swelling of the pattern due to cell remodeling, as opposed to intrinsic properties of the gel, as we did not see statistically significant differences in pattern swelling after 7 days of incubation in full serum media as compared to PBS in acellular gels (**Supplementary Figure S5**).

### Time-dependent softening of the matrix controls hMSC spreading

We next hypothesized that softening at different timepoints throughout the culture period would enable temporal control over cell spreading. To test this hypothesis, we encapsulated hMSCs in the stiffest DNs (5 mM PEG-NB: 20 mM dicysteine) and single network SPAAC gels. A subset of the DNs were softened on day 3 or day 5; all gels were fixed on day 7 (**Figure 5a**). Unlike in the patterned substrates, we observed that spreading area and eccentricity in SPAAC gels as compared to the DNs were not statistically significant (**Figures 5b-d**), with cells in SPAAC gels exhibiting slightly larger spread areas (385 ± 80 µm^2^) compared to those in the DNs (347 ± 43 µm^2^) with only slightly more branched morphologies. We noted that hMSCs proliferated more in the soft networks as compared to the other conditions (**Figure 5b**), potentially limiting their spreading time. Upon softening the DNs, hMSC morphology became branched, with cells exhibiting very prominent protrusions. Surprisingly, cells cultured in a stiff gel for 5 days displayed significantly higher spread areas (570 ± 47 µm^2^) as compared to SPAAC gels and DNs, but not compared to Day 3 (442 ± 45 µm^2^) (**Figures 5b-c**), and significant increases in eccentricity as compared to DNs (**Figure 5d**). We additionally saw very similar behavior in 10T1/2 fibroblasts (**Supplementary Figure S6**), with longer durations in a stiff matrix prior to softening eliciting robust spreading. This delayed spreading response could be due in part to increased cell-generated stresses in the stiff matrices, that upon softening, may result in increased matrix displacement and protrusion formation.^[55,56]^

**Figure 5.**
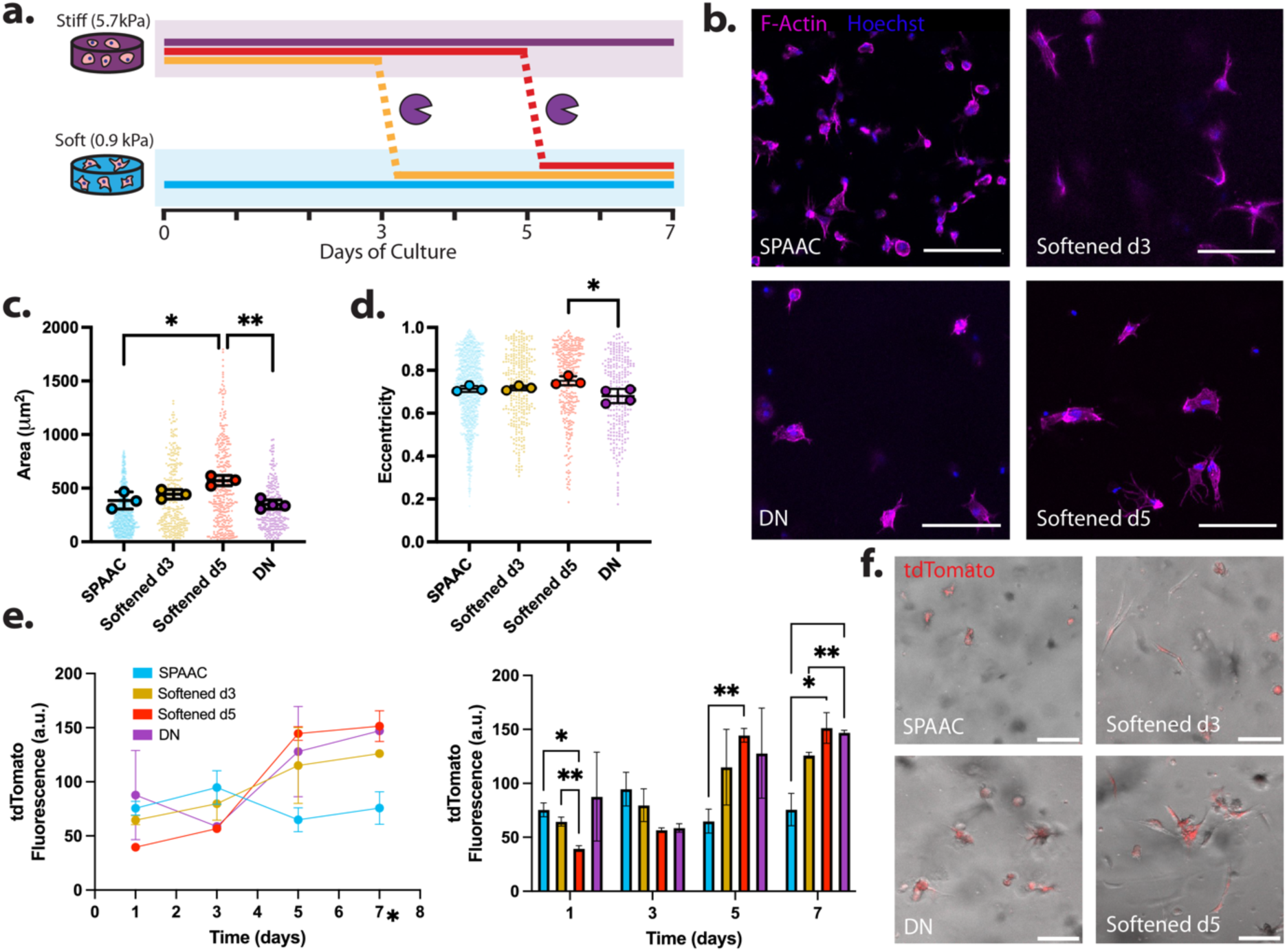
Time-dependent softening controls hMSC spreading behavior. **(a)** Experimental set up. hMSCs were encapsulated in static or dynamic DNs, or SPAAC networks. Dynamic DNs underwent softening on day 3 (“softened d3”) or day 5 (“softened d5”). **(b)** Actin and nuclear staining reveals distinct morphologies amidst experimental groups. Scale bar = 100 µm. **(c)** Quantification of cell area. Small dots indicate individual cell values, whereas larger circles indicate per gel average. Statistics were conducted on the per-gel averages. One-Way ANOVA, Tukey’s post-hoc test, *p = 0.0105, **p = 0.0021. **(d)** Quantification of eccentricity. One-Way ANOVA, Tukey’s post-hoc test, *p = 0.0123. **(e)** Live imaging of tdTomato intensity as a function of time in line (left) and bar (right) plot form. Dynamic DN gels were imaged prior to degradation on their respective degradation days. Two-Way ANOVA, Tukey’s post-hoc test. *p < 0.05, **p < 0.01 **(f)** Representative images of hMSCs expressing tdTomato upon TEAD binding events occurring on day 7 of encapsulation. Scale bar = 100 µm.

Given the previous results, we questioned whether this surprising spreading phenomenon could be due to differential translocation of YAP as part of the Hippo signaling pathway. To that end, we designed a tdTomato hMSC reporter line based on the Signalome reporter system for YAP/TAZ, which gives a live output of YAP translocation, but does not introduce exogenous YAP that may disrupt signaling processes.^[57]^ In this system, as the YAP/TAZ complex translocates to the nucleus and binds to the transcription factor TEAD, which then drives the expression of tdTomato (**Supplementary Figure S7a-b**).^[58]^ We repeated the dynamic softening experiment with this modified cell line and live-imaged each gel on days 1, 3, 5, and 7; gels softened on days 3 and 5 were first imaged on their respective days of softening, and then treated with 4S9. Initially, cells displayed a low level of tdTomato fluorescence, which began to increase with time in the DN conditions, with a sharp increase between days 3 and 5 (**Figures 5e-f**). By day 7 of culture, dynamic gels softened on day 3 saw a small drop in tdTomato intensity, but did not return to the levels of cells cultured in soft SPAAC gels; in a more pronounced fashion, the gels softened on day 5 exhibited the highest TEAD activity on day 7, yet not statistically significant from DN gels, with accordingly large spread areas (**Figure 5f, Supplementary Figure S7c**). The elevated Hippo signaling in static stiff and dynamic gels in comparison to static soft gels may signify a conserved memory of the previous stiff environment and may help facilitate the formation of advanced protrusions. While YAP translocation has been shown to be fast (on the order of minutes),^[59,60]^ downstream signaling and cytoskeletal reorganization in 3D may occur on much longer scales. Additionally, previous reports have similarly found that conditioning cells longer on a stiff matrix induces irreversible YAP activation; in these studies, 7 and 10 days of culture were found to induce an irreversible condition, whereas 1 day was reversible.^[61]^ In our studies, 5 days of stiff 3D culture similarly induces this retention of YAP in the nucleus; however future studies extending the time of soft culture could be useful in further probing this mechanical memory.

### Transcriptomic analysis of Caco-2 cells subjected to time-dependent softening

An added and significant advantage of our dynamically softening bulk hydrogels is that encapsulated cells can be recovered from the gels in a near “biologically invisible” manner and subsequently processed via single-cell methodologies. Towards this end, we sought to exploit the utility of our system to investigate cellular mechanomemory through global transcriptome quantification using RNAseq. One area where the concept of mechanomemory has recently gained traction is in cancer therapies;^[18]^ in particular, there has been growing interest in examining how modulating mechanics of the tumor microenvironment via antifibrotic treatments could improve outcomes in colorectal and pancreatic ductal adenocarcinoma.^[17,62,63]^ Of note, Shen et al. reported that altering the mechanical properties of metastatic colorectal tumor stroma by inhibiting fibroblast matrix deposition, improved patient response to targeted therapies;^[17]^ yet, a full mechanistic explanation remains elusive. Moreover, studies in hMSCs and gastric cancer cell lines have shown that cells retain memory of their previous mechanical environments, and that dosage time greatly affects regulation of YAP and downstream signaling pathways.^[6,25,61]^ Thus, we asked whether mechanomemory of colorectal cancer could be assayed with our system.

The colorectal adenocarcinoma cell line Caco-2 was selected as a model cell line for its popularity as a model for human intestinal mucosa^[64]^ and as a well-differentiated tumor model.^[65]^ We encapsulated Caco-2 cells in the baseline DN formulation (3 mM PEG-NB: 12 mM dicysteine: 3 mM PEG-BCN: 12 mM diazide) which matches this stiffness of primary, premetastatic colon cancer tissue collected from patients, as well as in soft, single network SPAAC gels, the moduli of which correspond to that of healthy colon tissue.^[46,66]^ A subset of the DN gels were softened, as in previous experiments, on day 3 and day 5. On day 7, all gels were fully degraded with a cocktail containing both 2A9 and 4S9 to mitigate potential effects of sortase treatment on genomic perturbation (**Figure 6a**). Collected cells were lysed for global transcriptome quantification by RNAseq, which identified and quantified over 15,000 genes from the human genome.

**Figure 6.**
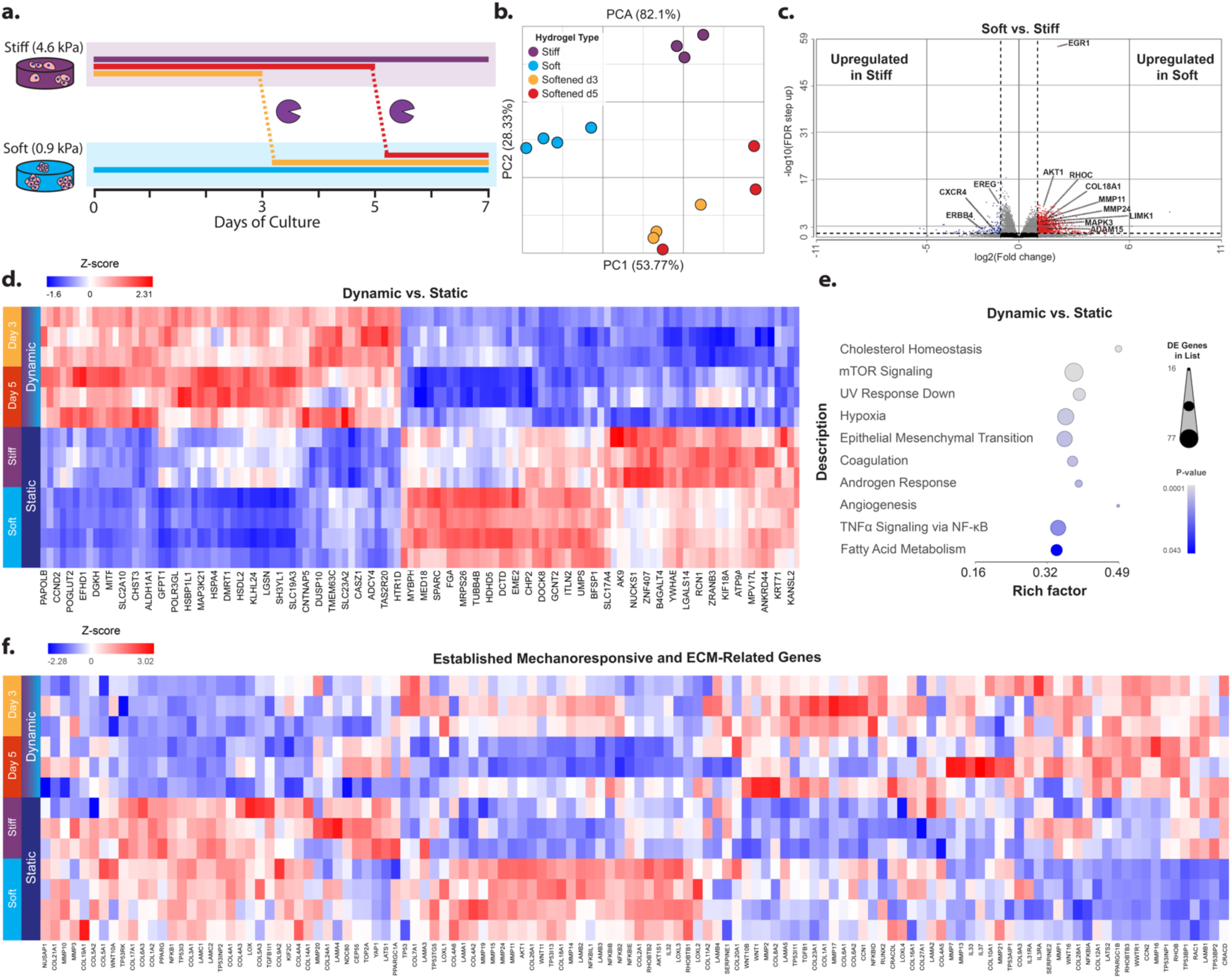
Transcriptomic analysis of Caco-2 cells in static and dynamically softened hydrogels. **(a)** Experimental set up. **(b)** PCA plot. **(c)** Volcano plot of gene expression profile in soft vs. stiff hydrogels. Red = upregulated genes in soft condition, blue = downregulated genes in soft condition. **(d)** Top upregulated and downregulated genes for dynamic vs. static comparison. **(e)** Enriched MSigDB gene sets among differentially expressed genes for dynamic vs. static comparison. **(f)** Heatmap of previously established mechanosensitive and ECM-related genes.

By principal component analysis (**Figure 6b**), the first principal component (PC) accounted for a relatively large fraction (54%) of sample variance and could distinguish readily between soft and stiff conditions. PC2 (28%) resolved the differences between static (soft and stiff) and dynamic (day 3 and day 5 conditions), however the day 3 and day 5 conditions clustered together, suggesting the day of softening is not as impactful as the bulk modulation of mechanics. To delve further into the various comparisons, we first chose to compare the two static, baseline controls—soft vs. stiff; we observed 1116 genes significantly upregulated and 131 downregulated (fold change > **|**2**|)** (**Figure 6c**). Over-representation analysis (ORA) via MSigDB identified terms related to cell division and metabolism, as well as MYC and mTOR signaling (**Supplementary Figure S8**). Certain previously established mechanosensitive genes such as *RHOC* and *LIMK1* were upregulated in the soft condition, suggesting higher cytoskeleton reorganization activity, which was also supported by upregulation of matrix and metalloproteinase genes, such as *COL18A1*, *COL4A2*, *MMP15, MMP24*, and *ADAM15* (**Figure 6c**).^[67–69]^ We also observed significant up and downregulation of genes up and downstream of the AKT1 and MAPK pathways, which are key regulators of proliferation, invasion, metabolism, cytokine production, and survival (e.g. *AKT1*, *MAPK3*, *EGR1*, *EREG*, *ERBB4, CXCR4*).^[70–73]^ Some of these genes have been shown to interact with the Hippo pathway; for instance, AKT1 has been demonstrated to phosphorylate YAP1, retaining it in the cytoplasm, and EREG and ERBB4, other genes upstream of the MAPK/AKT pathways, may also direct YAP1 localization to the nucleus.^[72,73]^ While many common mechanosensitive genes in the Hippo pathway did not exhibit high fold changes in expression, *YAP1*, *ROCK1,* and *ROCK2* expression were significantly lower in the soft conditions (FDR < 0.05, fold change -1.44, -1.51, -1.53, respectively), as has been previously shown in other biomaterial platforms, both 2D and 3D.^[6,61]^

When we compared dynamic (day 3 and day 5) vs. static (stiff and soft) conditions, we saw striking differences in gene expression (**Figure 6d**). ORA of the differentially expressed genes returned pathways such as “cholesterol homeostasis”, “mTOR signaling”, “hypoxia” and “EMT” as highly significant (**Figure 6e**). Similarly, we conducted gene-set enrichment analysis (GSEA),^[74]^ which takes into account all differences in gene expression even below the threshold required for ORA, on the same comparison, and saw downregulation of cholesterol homeostasis and EMT-related genes in dynamic conditions, with similar trends in glycolysis genes (**Supplementary Figure S9**). We then conducted ORA on differentially expressed genes in all comparisons (**Supplementary Figure S8**). While the comparison for day 3 vs. day 5 did not yield any significant pathways, all other comparisons showed enrichment of many similar pathways: mTOR and MYC-related signaling pathways were highly enriched in 5 out of 5 and 4 out of 5 comparisons, respectively. We observed striking differences in these signaling pathways, especially between the dynamic conditions compared to stiff or soft (**Supplementary Figure S10**). Given that both these pathways are downstream of PI3K-AKT and MAPK activity, it is unsurprising that they are highly affected by mechanical stimuli, and have previously been demonstrated as such.^[75–78]^ The protooncogenic protein MYC has been demonstrated to regulate global metabolic reprogramming and plays a pivotal role in 5-fluorouracil resistance in colorectal cancer.^[79,80]^ Many MYC-associated genes were most highly expressed in the stiff condition, while the dynamic conditions displayed the lowest expression of these genes (**Supplementary Figure S10**), similar to a recent report by Nguyen and Lin in COLO-357 spheroids,^[81]^ potentially signifying that modulating the mechanical environment plays a major role in suppressing this signaling axis.

Only 147 genes were differentially expressed between day 3 and day 5, with 9 being either significantly upregulated or downregulated (FDR < 0.05, |fold change| ≥ 2; 8 up, 1 down). Intriguingly, two ECM-related proteins *COL18A1* and vinexin (*SORBS3*) were downregulated in stiff and day 5 conditions. This, and our other results, demonstrate that while the day of softening is not as powerful in eliciting a strong differential response, bulk mechanical manipulation of the 3D environment produces large-scale signaling responses.

We additionally examined how these changes in mechanical properties affect known mechanosensitive and ECM-related genes previously shown in 3D biomaterial platforms (**Figure 6f**).^[25]^ We noted differences in gene expression among static and dynamic conditions, as well as differences between the static soft and stiff conditions. Further, we noticed a gradient-like response (middle and right side of heatmap) for a subset of MMP (specifically many membrane-bound subtypes) and collagen and laminin genes, with soft gels having the highest expression, followed by intermediate expression in the day 3 condition, and lower expression in the day 5 condition, similarly to the stiff condition. ECM remodeling and deposition are complex and multistep processes, known to take place on longer timescales than acute intracellular signaling cascades, potentially explaining the range in response across multiple days.^[82–84]^ In sum, 4D-tunable biomaterials, such as our DN system, are key to exploring these intricate, interwoven, and dynamic biological phenomena. Our DN design is the first fully synthetic interpenetrating network system that enables reversible and spatially controlled mechanical modulation with subsequent bioorthogonal cell recovery. The combination of orthogonal single network formation and degradation reactions opens powerful doors for investigating the role of matrix mechanics on many biological systems.

## Conclusion

3D models that can modulate native tissue mechanics in a spatiotemporally defined manner could prove useful in studying the effects of ECM stiffness on drug efficacy, disease progression, and mechanomemory.^[18,23]^ Herein, we have presented a bioorthogonal platform that allows for spatiotemporal control over Young’s moduli, as well as temporal control over bulk stiffening/softening and subsequent cellular release. We engineered the stepwise formation of a DN composed of two discrete bioconjugate chemistries which can then be reversibly stiffened through distinct bioorthgoonal reactions and subsequently softened via orthogonal sortase treatments. In our design, all macromer components can be either initially present, or sequentially diffused in: the SPAAC network forms spontaneously, generating a soft initial network, that can then be stiffened with spatial precision by thiol-ene photopolymerization. The DN can be subsequently reverted back to the starting single network via sortase-mediated transpeptidation; the photopolymerization and enzymatic degradation can be repeated as one sees fit, but then the whole network can be fully degraded to recover embedded cells in a “biologically invisible” manner – a feature not common in many current biomaterial models – for downstream analysis.

In developing this system, we demonstrate its utility towards studying changes in cell morphology and mechanosignaling, both through live imaging and bulk transcriptomic analysis. Moreover, our material chemistry-based approach is generalizable: other bioorthogonal chemistries and/or enzymatic pairs may be utilized to create alternate versions of reversibly formed DNs depending on access and need. We readily envision the extension of our method to other biological systems where mechanobiological questions are of pressing interest.

## Acknowledgements

This work was supported by a Faculty Early Career Development (CAREER) Award (DMR 1652141 to C.A.D.), a standard award (DMR 1807398, C.A.D.), and Graduate Research Fellowships (DGE 1762114 to I.K. and R.C.B.) from the National Science Foundation; a Maximizing Investigators’ Research Award (R35GM138036 to C.A.D.) and an Interdisciplinary Training Grant (T32CA080416 to I.K.) from the National Institutes of Health. M.C.R. and work conducted in the Institute for Stem Cell and Regenerative Medicine (ISCRM) Genomics Core were supported by a generous gift from the John H. Tietze Foundation. We also acknowledge ISCRM (research scholarship to R.C.B.), the Mary Gates Endowment for Students (research scholarship to E.C.G.), and the Louis Stokes Alliance for Minority Participation (LSAMP) Program (research scholarship to K.R.V). The authors would additionally like to acknowledge Dr. Micah Glaz at the UW Molecular Analysis Facility for assisting with AFM method development, Dale Whittington and Dr. J. Scott Edgar (in memoriam) for assisting with LCMS, and Xinting Li for peptide synthesis help.

## Author Contributions

I.K., E.C.G., R.C.B., and C.A.D. conceived and designed the experiments. I.K., E.C.G, K.R.V., and R.C.B. executed synthesis of material precursors, protein expression, and mass spectrometry. K.R.V. conducted photorheology experiments. I.K. conducted AFM, fluorescent release, and encapsulated cell culture experiments. E.C.G. carried out and analyzed cellular patterning experiments. S.Y. conducted multiphoton patterning and imaging. I.K. and J.W.H. designed and created the hMSC TEAD-reporter line. I.K., M.C.R., and J.W.H. conducted RNAseq library prep, sequencing, and analysis of data. I.K. and C.A.D. wrote the manuscript. I.K. and C.A.D. funded the experiments. C.A.D. provided mentorship and lab space.

## Data Availability Statement

The data that support the findings of this study are available from the corresponding author upon reasonable request.

