## Supplementary Information for "Stepwise Stiffening/Softening of and Cell Recovery from Reversibly Formulated Hydrogel Double Networks"

**Method S1 – Macromer Synthesis**

4-arm poly(ethylene glycol)-bicyclononyne (PEG-BCN) was synthesized as previously described.[1] 4-arm poly(ethylene glycol) tetraamine (MW 20kDa, 1.143 g, 0.0571 mmol, 0.2 mol $NH_2$ groups, 1x; JenKem Technology USA; Plano, TX) and (1R,8S,9s)-bicyclo[6.1.0]non-4-yn-9-ylmethyl (2,5-dioxopyrrolidin-1-yl) carbonate (BCN-OSu) (100 mg, 0.343 mmol, 1.5x to $NH_2$ groups) were dissolved in anhydrous dimethylformamide (5 mL) and N,N-Diisopropylethylamine (DIEA, 159µL, 8 mmol, 4x to $NH_2$ groups) and stirred overnight. The following day, the mixture was diluted with water (5x volume) and dialyzed in DI $H_2O$ overnight (molecular weight cutoff ~ 2kDa; SpectraPor, Repligen; Waltham, MA), lyophilized to yield a white powder, and resuspended in PBS at a 10 mM stock concentration. $^1$H NMR confirmed functionalization to be >95% by comparing integral values for characteristic BCN peaks (δ 2.24, 1.57, 1.34, 0.92) with those from the PEG backbone (δ 3.63).

8-arm poly(ethylene glycol)-norbornene (PEG-NB) was synthesized as described previously.[2] 5-norbornene-2-carboxylic acid (196 µL, 1.6 mmol, 8x) was prereacted for five minutes under $N_2$ with 2-(1H-7-Azabenzotriazol-1-yl)-1,1,3,3-tetramethyl uranium hexafluorophosphate methanaminium (HATU) (601 mgs, 1.58 mmol, 3.95x) and DIEA (557 µL, 3.2 mmol, 8x) in 5 mLs of anhydrous DMF, then transferred to the flask containing 8-arm PEG amine (pentaerithrytol core) (MW 20kDa, 1 g, 0.05 mmol, 0.4 mol $NH_2$ groups, 1x; JenKem Technology USA; Plano, TX) and reacted overnight under $N_2$. The following day, PEG-NB was precipitated in ice-cold diethyl ether, resuspended in DI $H_2O$, and dialyzed overnight. Afterwards, it was lyophilized to yield a white powder and resuspended in PBS at a 30 mM stock concentration. Norbornene conjugation was assessed with $^1$H NMR (500 MHz, $CDCl_3$) by comparing the characteristic norbornene peaks (δ = 5.9 – 6.3 ppm) to those of the PEG backbone (δ 3.63 ppm).

**Method S2 – Peptide Synthesis**

The MMP- AND 2A9-sensitive crosslinker H-RGPQGIWGQLAETGGRK(dde)-$NH_2$ was synthesized on rink amide ProTide resin (CEM Corporation; Charlotte, NC) following standard microwave-assisted Fmoc solid phase techniques with HBTU activation (CEM Liberty 1; Charlotte, NC) at a 0.25 mmol scale. The 1-(4,4-dimethyl-2,6-dioxacyclohexylidene)ethyl (dde) group was deprotected by treating with 2% hydrazine monohydrate in dimethylformamide (DMF, 3x10 min). The N-terminal amine as well as the lysine ε-amino group of the C-terminal lysine were simultaneously coupled with 4-azidobutanoic acid (227 µL, 2 mmol, 4x to $NH_2$ groups) via HATU activation (750 mg, 1.97 mmol, 3.95x to $NH_2$ groups) and DIEA (1.38 mL, 8 mmol, 16x to $NH_2$ groups) and reacted for 1.5 hours. Reaction efficiency was tested via the Kaiser reaction with amines. The resin was treated with trifluoracetic acid (TFA)/triisopropylsilane (TIS)/water (95:2.5:2.5) for 3 hours, then crashed out and washed in ice-cold diethyl ether (150mL). The crude peptide was purified via semi-preparative reversed-phase high performance liquid chromatography with a linear gradient of 5-100% acetonitrile and 0.1% TFA for 45 minutes and then lyophilized to yield a white powder of the final peptide $N_3$-RGPQGIWGQLAETGGRK($N_3$)-$NH_2$. Peptide mass was verified via ESI-LCMS. The peptide was aliquoted at a 50 mM stock concentration [20% DMSO in phosphate buffered saline (PBS), pH = 7].

The fibronectin-derived adhesion peptide H-GRGDS-$NH_2$ (RGDS) was synthesized and purified in the same manner as above, reacted with 4-azidobutanoic acid to yield the N-terminally monofunctionalized peptide $N_3$-GRGDS-$NH_2$, and aliquoted at a 50 mM stock concentration. The H-CRGDS-$NH_2$ peptide was similarly synthesized, but cleaved with a TFA/TIS/ethanedithiol/water (94:2.5:2.5:1) cocktail. It was similarly purified, lyophilized, and then resuspended in 10% glacial acetic acid at a known concentration, and lyophilized a second time for storage.

**Method S3 –2A9 and 4S9 Expression and Purification**

Electrically competent BL21 cells were transfected with the respective eSrtA plasmids and selected on kanamycin-containing agar plates. 5 mL of Luria Broth (LB) with kanamycin (50µg/mL) was inoculated with a plasmid-containing colony and allowed to grow overnight at 37ºC at 200 rpm. The following day, 1 L of LB broth with kanamycin was inoculated with the 5mL overnight culture and allowed to incubate at 37ºC, 200 rpm until the O.D. reached 0.6, at which point isopropyl β-d-1-thiogalactopyranoside (IPTG) was added to the culture at a final concentration of 0.5mM to induce protein expression. The culture was then moved to 18ºC, 200 rpm and allowed to incubate overnight. Cells were pelleted via centrifugation (4000g, 20 mins), resuspended in lysis buffer (20 mM Tris, 50 mM NaCl, 10 mM imidazole; pH 7.5) with 1 mM phenylmethylsulfonyl (PMSF; TCI, Portland, OR) protease inhibitor, and lysed with sonication (6x, 3 min cycle, 30% amplitude; Fisher Scientific; Waltham, MA). The lysate was clarified via centrifugation (20 mins, 7500g) and filtered with a 0.45 µm syringe filter (Sartorius; Göttingen, DE). The clarified lysate was then purified using an ÄKTA Pure 25 L FPLC (Cytiva; Marlborough, MA) equipped with a 5 mL HisTrap HP column at a flow rate of 5 mL $min^{-1}$. The column was equilibrated with 5 column volumes of lysis buffer, followed by loading the sample and washing with 8 column volumes of endotoxin removal buffer (20 mM Tris, 50 mM NaCl, 20 mM imidazole, 0.1% Triton-X 114; pH = 7.5) and 8 column volumes of wash buffer (20 mM Tris, 50 mM NaCl, 20 mM imidazole; pH = 7.5). Triton-X 114 removal was monitored by ultraviolet absorbance (λ = 280 nm). His-tagged protein was eluted over an 8-column volume gradient of imidazole (20 mM Tris, 50 mM NaCl, 5-250 mM imidazole; pH = 7.5) into a 96 well plate; protein-containing fractions were pooled and dialyzed into PBS over two days. Purified sortase was spin concentrated with an Amicon Ultra 15 centrifugation filter (10,000 Da cut off) and diluted with PBS containing 20% glycerol to a final concentration of 100 µM (2x) and 200 µM (4X). Sortase purity was evaluated with sodium-dodecyl sulfate-polyacrylamide gel electrophoresis (SDS-PAGE) and identity was confirmed with electrospray ionization mass spectrometry on an AB SCIEX 5600 QTOF instrument (SCIEX; Framingham, MA).

**Method S4 – Measuring Gel Swelling and Water Content**

SPAAC gels were cast at concentrations of 3 mM PEG-BCN: 6 mM diazide crosslinker. DN gels were cast at concentrations of 3 mM PEG-BCN: 6 mM diazide crosslinker: 3 mM PEG-NB: 12 mM dicysteine crosslinker. Dynamic gels were formed as DNs, allowed to swell in PBS for 4 hours, softened with 4S9, and then washed 3 x 10 mins in water to wash out residual thiol-ene components. Gels were allowed to swell overnight in DI $H_2O$, then gently dabbed to remove excess water, and weighed. The gels were then allowed to dry overnight and once again, weighed. Gel swelling ratio (Q) and water content were calculated according to the following equations:[3]

$$\boldsymbol{Q} = \frac{M_{gel}}{M_{polymer}}$$

$$\boldsymbol{Water\ content}\ (\%) = 100\% \times \frac{M_{gel} - M_{polymer}}{M_{gel}}$$

where $M_{gel}$ is the mass of the swollen hydrogel and $M_{polymer}$ is the mass of the dried gel.

**Figure S1 – Electrospray ionization liquid chromatography mass spectrometry (ESI-MS) of 2A9-responsive diazide peptide crosslinker and $N_3$-GRGDS.**

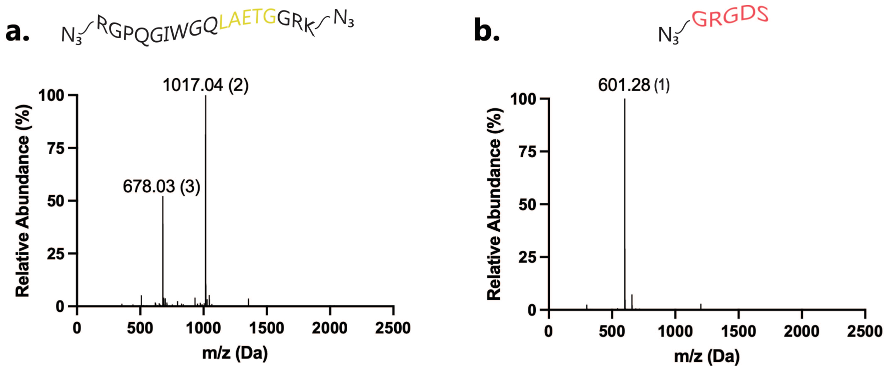

Expected masses ($N_3$-GRGDS-$NH_2$: 600.58 g/mol and $N_3$-RGPQGIWGQLAETGGRK($N_3$)-$NH_2$: 2034.06 g/mol) were the dominant observed peaks. The charge number of ions is indicated with parentheses.

**Figure S2 – SDS PAGE and ESI-MS of 4S9 and 2A9**

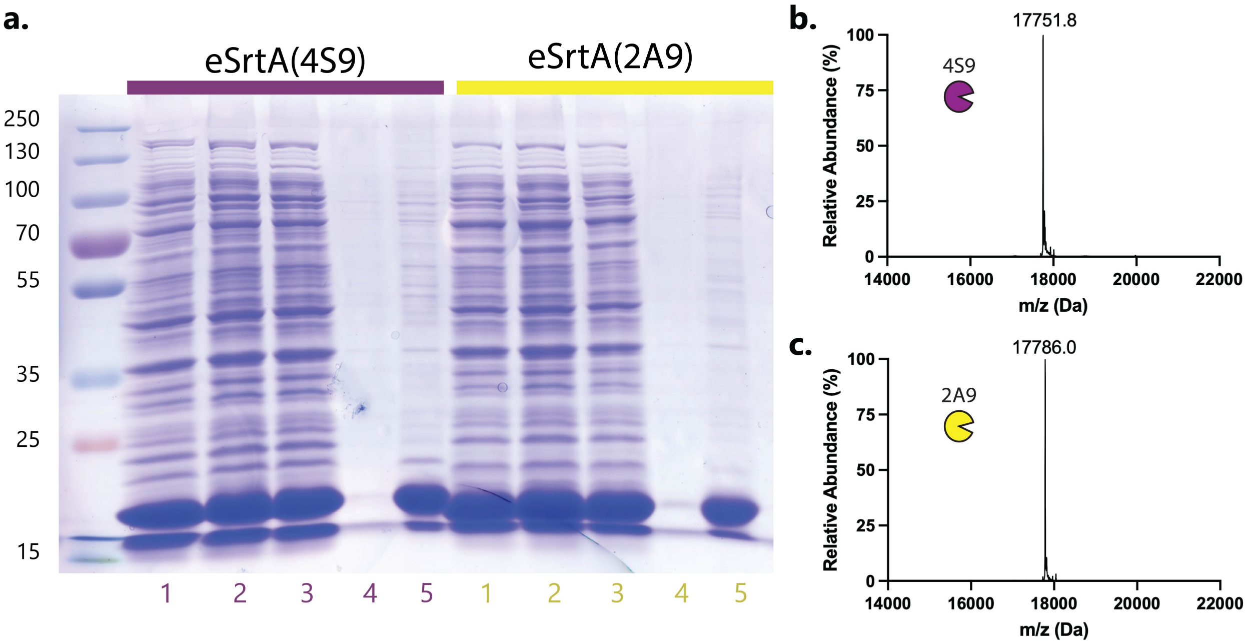

SDS-PAGE and ESI-MS were used to evaluate the purity of 4S9 and 2A9. **(a)** Strong bands at around 18 kDa in both final concentrated samples were observed with SDS-PAGE. (1) Clarified lysate, (2) flow-through, (3) wash 1, (4) wash 16, (5) concentrated sample. **(b)** ESI-MS analysis of 4S9. Expected mass: 17,751.97 g/mol, observed mass: 17,751.8 g/mol. **(c)** ESI-MS analysis of 2A9. Expected mass: 17,785.87 g/mol, observed mass: 17,786.0 g/mol.

**Figure S3 – 3D CAD Model for Two-Photon Thiol-Ene Polymerization**

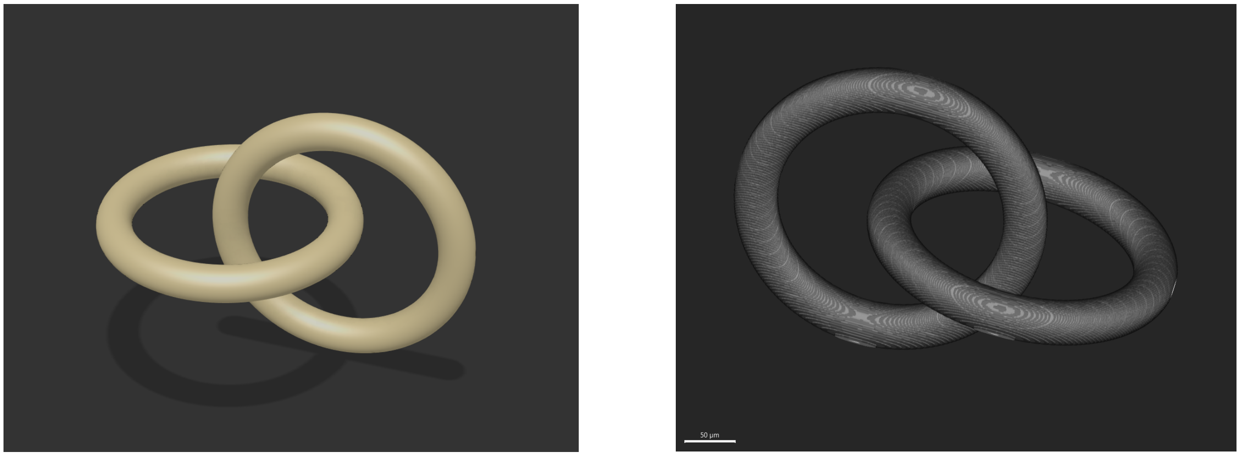

A 3D model of intersecting rings was constructed using Autodesk Fusion 360. The model was then sliced in Autodesk Netfabb Premium 2021 along the z-direction into a 235-image sequence (1024 px × 1024 px) with defined voxel size of 0.44 µm × 0.44 µm × 1 µm (L × W × H), which was then loaded into ScanImage (MBF Bioscience; Williston, VT) as the reference image stack for z-stack image-guided multiphoton patterning. Scale bar = 50 µm.

**Figure S4 – Influence of RGDS on Young's Moduli of Bulk Gels**

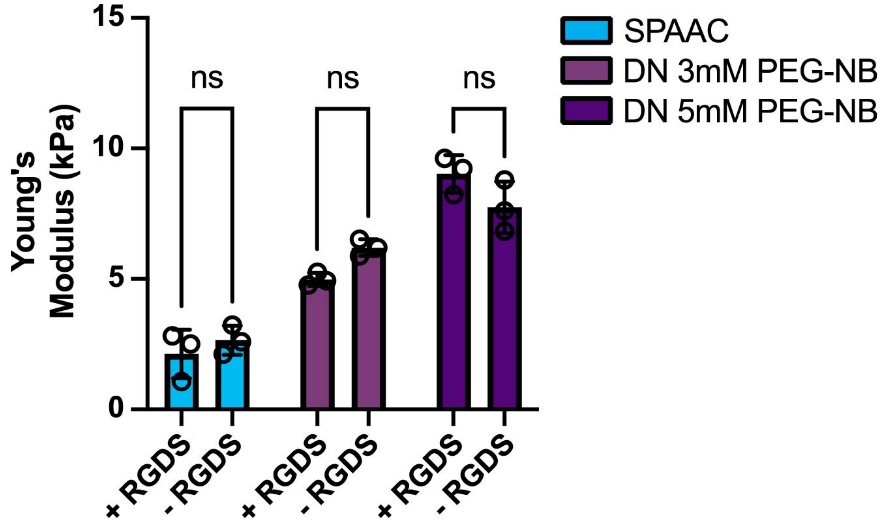

AFM measurements showed slightly lower, but not statistically significant, Young's moduli in gels modified with RGDS, except in the stiffest DN formulation (Two-way ANOVA, Tukey's Post-hoc test). SPAAC gels were cast at concentrations of 3 mM PEG-BCN: 6 mM diazide crosslinker ± 1 mM RGD. DN gels were cast at concentrations of 3 mM PEG-BCN: 6 mM diazide crosslinker: 3 mM PEG-NB: 12 mM dicysteine crosslinker ± 1 mM RGDS.

**Figure S5 – Swelling Quantification of Bulk and Patterned Gels**

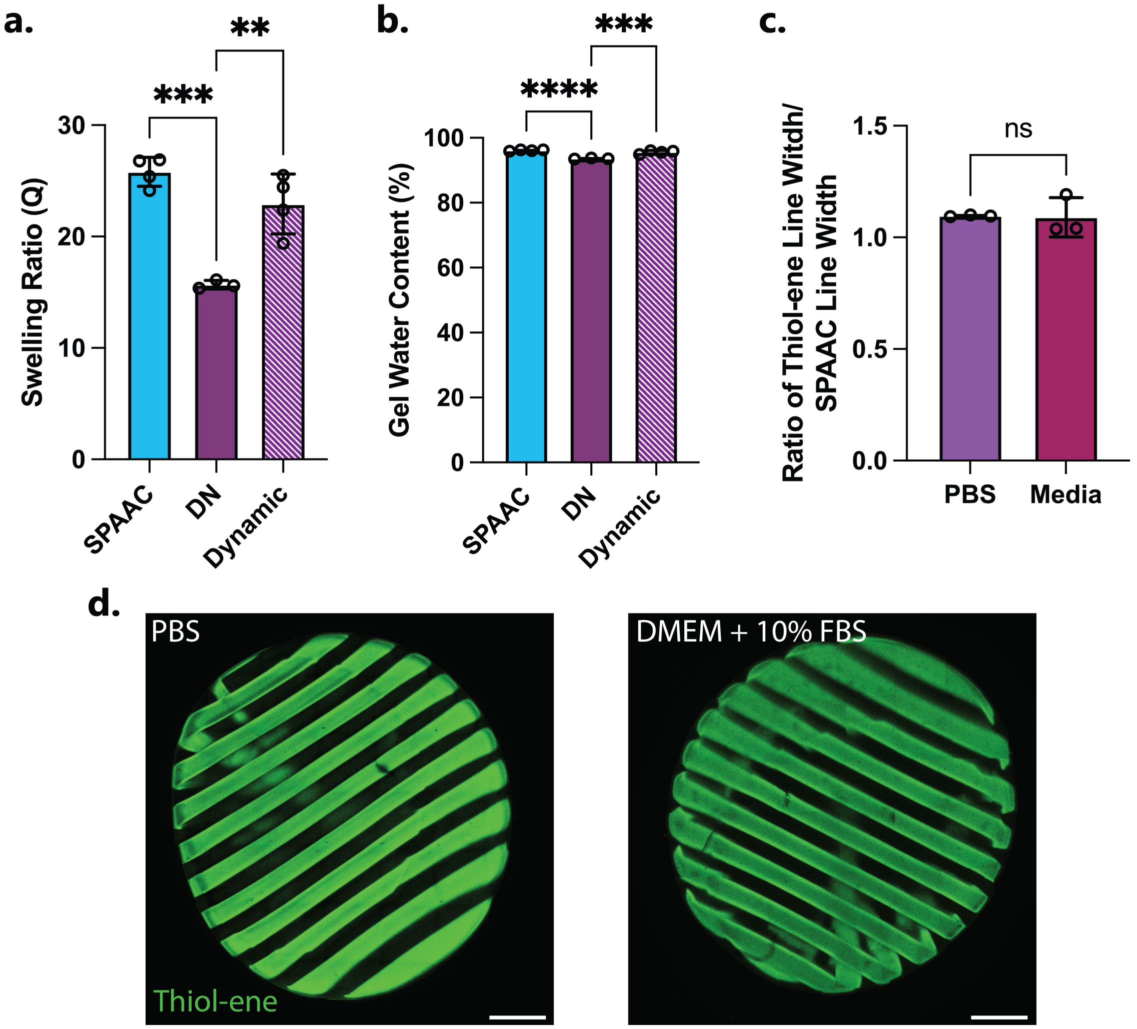

Swelling ratios **(a)** and gel water content **(b)** were calculated as described in the SI methods. SPAAC gels were cast at concentrations of 3 mM PEG-BCN: 6 mM diazide crosslinker. DN gels were cast at concentrations of 3 mM PEG-BCN: 6 mM diazide crosslinker: 3 mM PEG-NB: 12 mM dicysteine crosslinker. Dynamic gels were softened with 4S9. Statistical differences for both parameters were noted between SPAAC and DN gels and Dynamic and DN gels (One-Way ANOVA, Tukey's Post-hoc test, $**p < 0.01$, $***p < 0.001$, $****p < 0.0001$). **(c)** 200 µm lines were lithographically patterned in gels using a chrome photomask. Ratios of the widths of the thiol-ene lines to the width of the SPAAC lines in patterned gels were measured in gels swelled overnight first in PBS and then swelled for 7 days in cell culture media. No significant differences were noted in pattern swelling. **(d-e)** Representative maximum intensity projections (z stack = 100 µm) of patterned gels swollen in PBS and DMEM. Scale bars = 1 mm.

**Figure S6 – Temporal Softening of DNs controls Fibroblast Spreading**

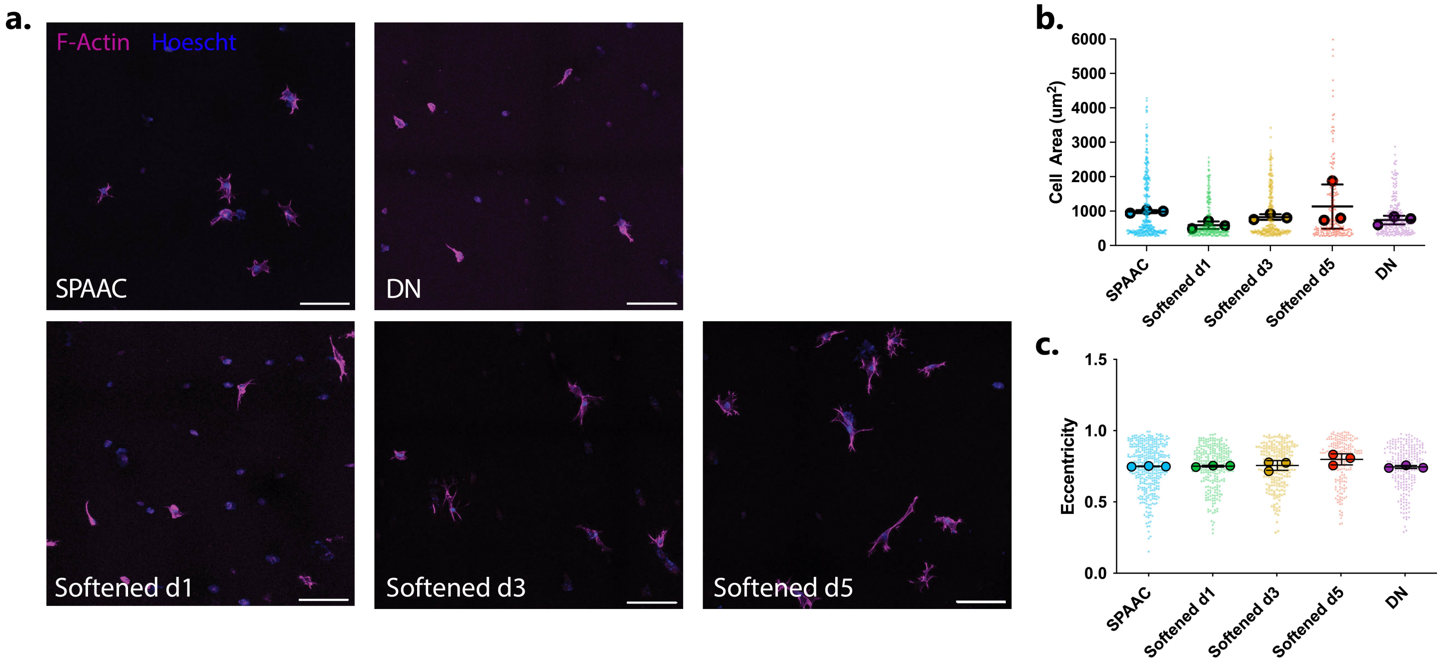

10T1/2 fibroblast spreading is dictated by day of softening. **(a)** Representative MIP images of encapsulated fibroblasts. Scale bar = 100 μm. **(b)** Quantification of cellular area. Significance was noted on the per cell level, similar to the hMSC experiments, but not on the per gel level. **(c)** Quantification of eccentricity.

**Figure S7 – tdTomato-TEAD hMSC Reporter Cell Line**

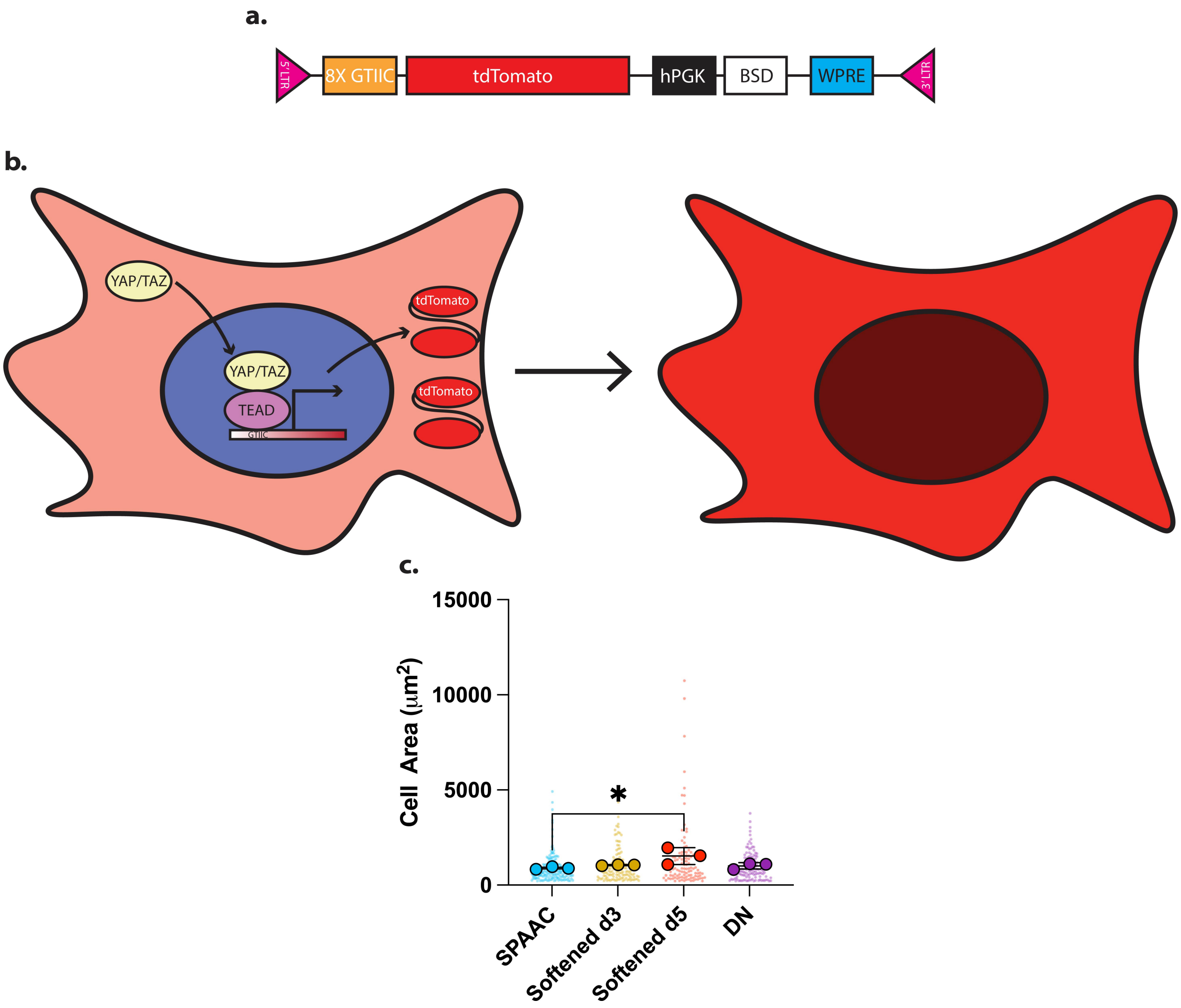

**(a)** tdTomato-TEAD reporter construct. GTIIC represents the TEAD-binding sequence. **(b)** Schematic of reporter cell line output. **(c)** Quantification of fixed cell area on day 7 of culture.

#### Figure S8 – Over Representation Analysis Plots of Individual Comparisons via MSigDB

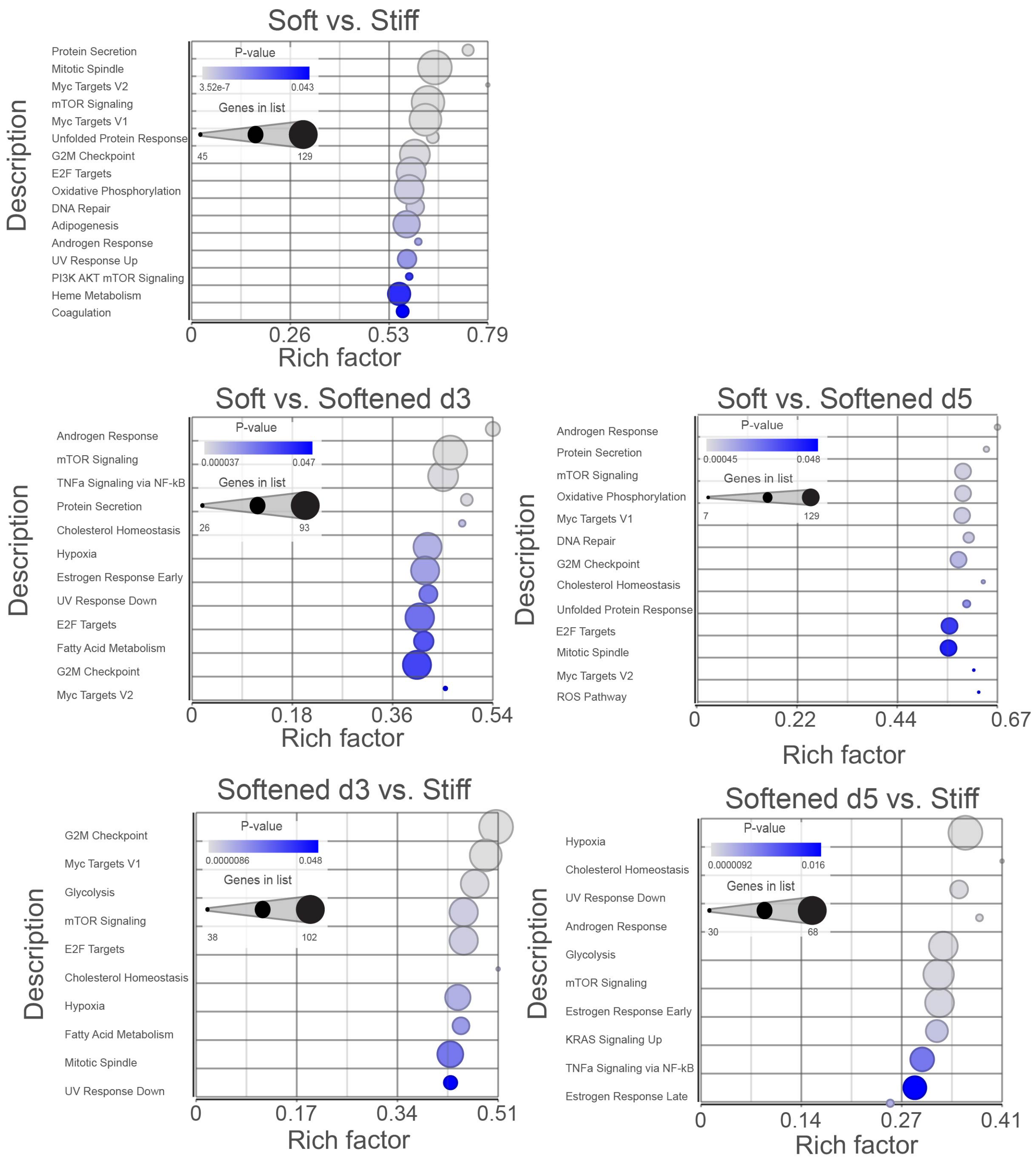

Overrepresentation analysis (ORA) was completed on all individual comparisons. Day 3 vs Day 5 comparisons did not yield any statistically significant ($p < 0.05$) pathways.

### Figure S9 – GSEA Plots

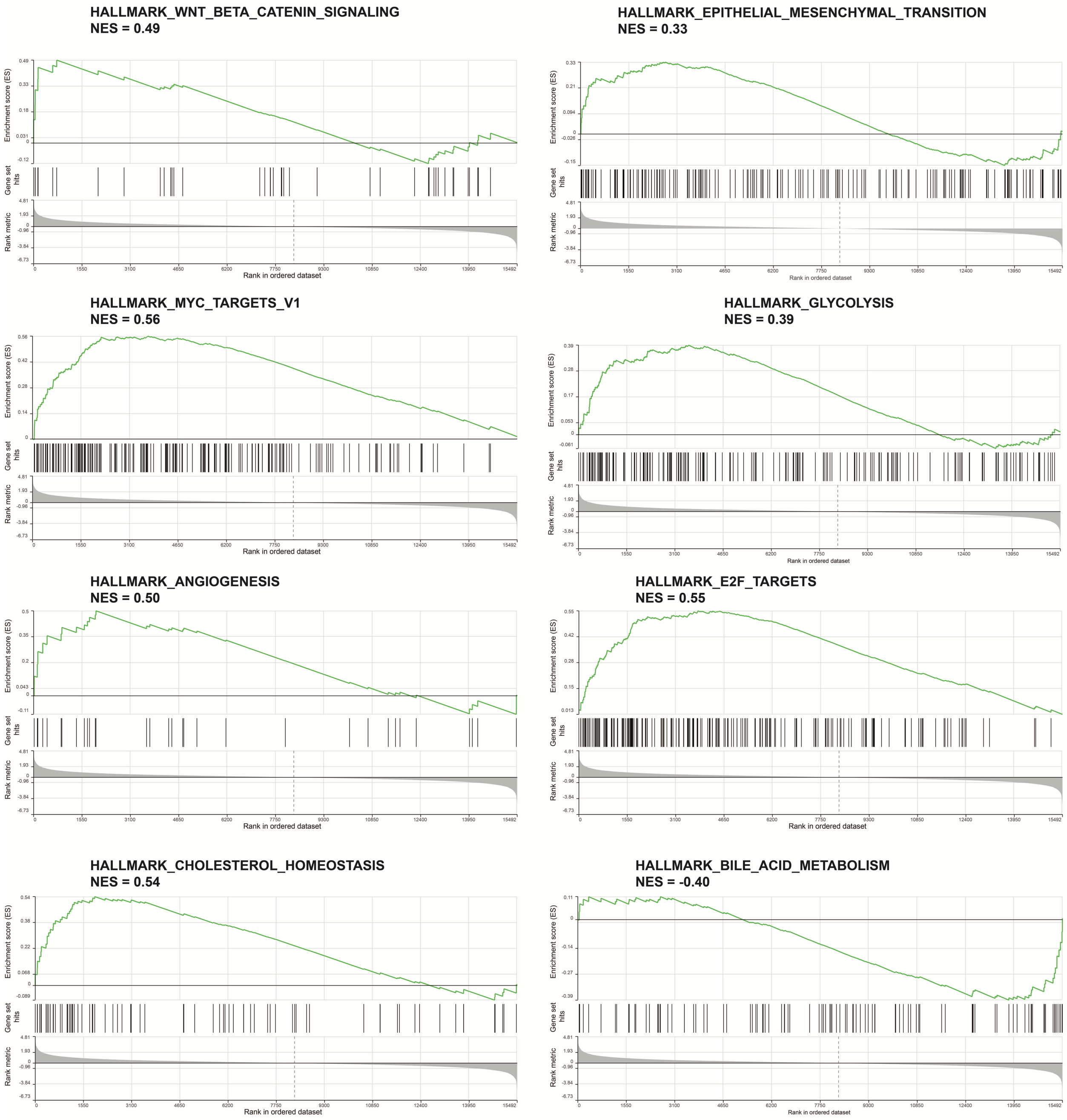

GSEA plots for pathways found to be significantly enriched in the static vs. dynamic comparison ($p < 0.05$). NES > 0 signifies upregulation in the static conditions, whereas NES < 0 signifies upregulation in the dynamic conditions.

**Figure S10 – Myc and mTOR Signaling Pathway Heatmaps**

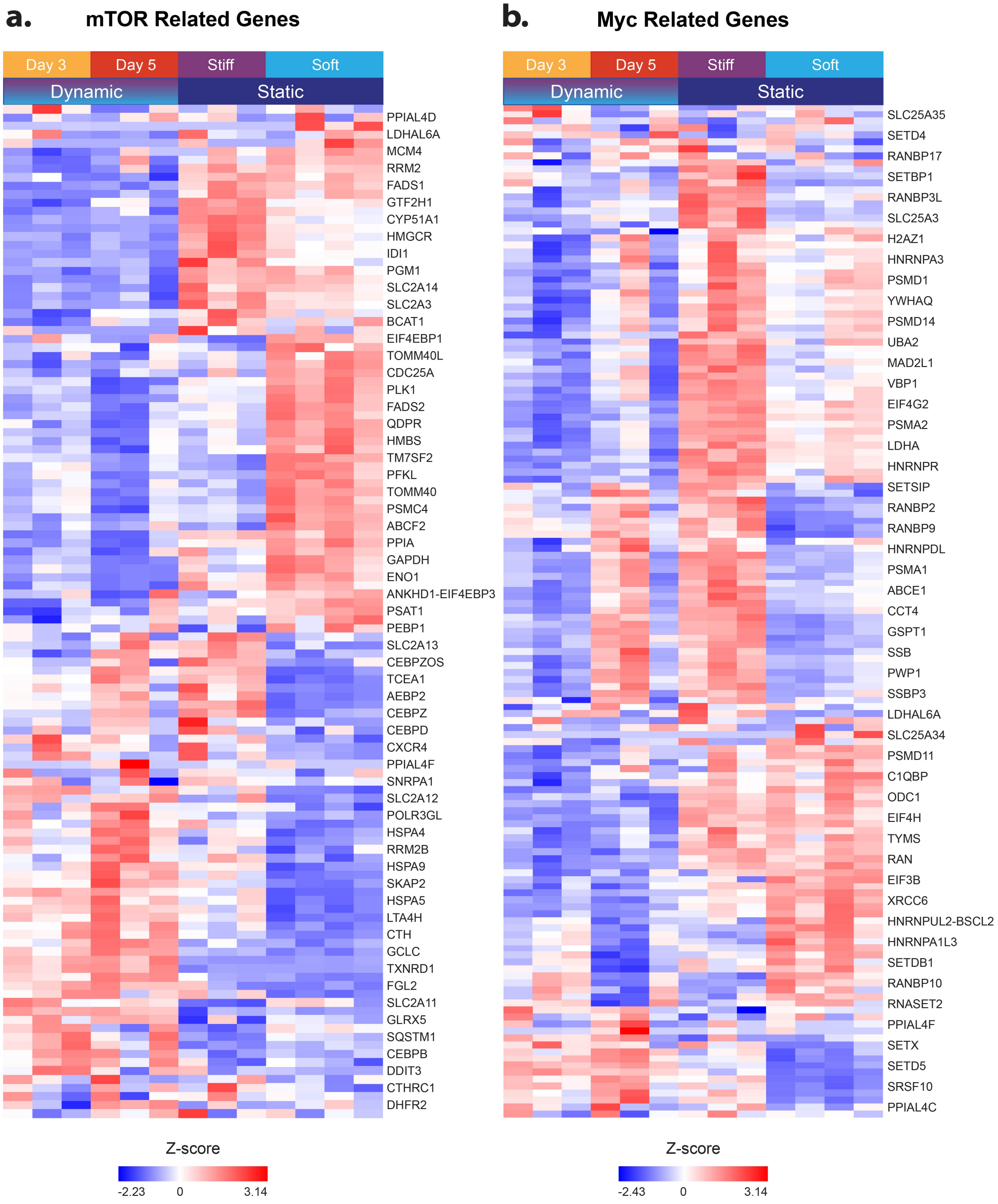

Expression profiles for genes related to **(a)** mTOR signaling and **(b)** Myc signaling as defined by MSigDB (MYC TARGETS V1 and V2).
